# *In vivo* Protein Footprinting Reveals the Dynamic Conformational Changes of Proteome of Multiple Tissues in Progressing Alzheimer’s Disease

**DOI:** 10.1101/2023.05.29.542496

**Authors:** Ahrum Son, Hyunsoo Kim, Jolene K. Diedrich, Casimir Bamberger, Daniel B. McClatchy, John R. Yates

## Abstract

Numerous studies have investigated changes in protein expression at the system level using proteomic mass spectrometry, but only recently have studies explored the structure of proteins at the proteome level. We developed covalent protein painting (CPP), a protein footprinting method that quantitatively labels exposed lysine, and have now extended the method to whole intact animals to measure surface accessibility as a surrogate of in vivo protein conformations. We investigated how protein structure and protein expression change as Alzheimer’s disease (AD) progresses by conducting in vivo whole animal labeling of AD mice. This allowed us to analyze broadly protein accessibility in various organs over the course of AD. We observed that structural changes of proteins related to ‘energy generation,’ ‘carbon metabolism,’ and ‘metal ion homeostasis’ preceded expression changes in the brain. We found that proteins in certain pathways undergoing structural changes were significantly co-regulated in the brain, kidney, muscle, and spleen.

## Introduction

The proper functioning of cellular machinery depends on the ability to maintain the functional structures of proteins. Proper folding of proteins is necessary to engage with partners in complexes and to perform catalytic activities. Protein folds or shapes can be measured by powerful, high-resolution *ex vivo* techniques such as X-ray crystallography, NMR, and Cyro-electron microscopy (Cyro-EM).^1–4^ Cyro-EM can be used to analyze large protein complexes if they are extracted from cells or are produced recombinantly prior to deposition on the grid and frozen.^5–7^ Multiplexed Ion Beam imaging (MIBI) and ion beam tomography are capable of imaging cells and tissues, but they are not explicitly used to study the structure of proteins and protein complexes. Modeling algorithms can generate protein structures from *ex-vivo* protein cross-linking data, while *in vivo* cross-linking analyses generate protein-protein interaction data. Thus, because no methods are available to determine the high-resolution structures of proteins *in vivo,* we are still limited in our ability to elucidate the structures of proteins in the cellular milieu.

Protein “footprinting” methods were developed to probe the folding and interactions of proteins (such as epitope sites in antigens) using protease restriction or covalent labeling to identify exposed regions of proteins.^8^ The data generated in protein footprinting experiments is often low resolution, but the potential scale of experiments has made it an attractive method. In 2010 West et al. showed proteome-scale footprinting in *S. cerevisiae* to determine off target binding of rapamycin^9^. A variety of protein labeling methods have been developed that provide low resolution *ex vivo* structural information about proteins.^5, 10–13^ Picotti and colleagues developed a limited proteolysis method to map ligand binding and protein folding in cell lysates and biofluids.^5, 14–16^ In 2015 Espino et al. used lasers to activate hydroxy radicals *in vivo* to label proteins, providing the first attempt to footprint an intact cell.^17^ Their approach has now been extended to the transparent worm *C. elegans*, which was chosen so the laser beam could penetrate the worm.^18^

Bamberger et al. developed Covalent Protein Painting (CPP), a chemical approach for quantitative protein footprinting to measure *in vivo* changes to protein conformations on a proteome scale.^19^ In CPP formaldehyde, a chemical that rapidly permeates through cells and tissues, is used to label proteins by forming a Schiff’s base at solvent exposed lysine residues. These unstable intermediates are converted to dimethyl labels by reduction with cyanoborohydride. After lysis of cells or tissue, denaturation and digestion of proteins, a second labeling with a different “weight” reagent is performed to label inaccessible amino acid residues. By using heavy and light isotope versions of the reagents, a quantitative measure of lysine accessibility can be obtained. Using this method, Bamberger *et al*. probed the conformational changes of a proteome from postmortem brain tissue to reveal structural changes and altered protein-protein interactions in the brain tissue of AD patients.^19^ In another study, Bamberger et al. measured the altered conformations of proteins in 60 cancer cell lines (NCI60).^20^ Because the CPP protein labeling method begins with the widely used formaldehyde fixation step for *in vivo* dimethyl labeling it should be extensible to whole animal labeling to study models of disease.

Methods to measure alterations of protein conformations *in vivo* are needed to study diseases caused by protein misfolding that create loss or gain of function disruptions to biological processes, including Alzheimer’s disease, a common misfolding disease that is characterized by plaques of amyloid-beta and tangles of tau proteins. As observed by Bamberger et al., late-stage neurodegenerative diseases in humans are characterized by the misfolding of many additional proteins, suggesting that there is a generalized failing of proteostasis.^19^ Techniques that allow *in vivo* measurement of protein folding would be a powerful tool for the study of these misfolding diseases.

Here, we used AD as a model to test our hypothesis that the global measurement of structural changes of proteins in tissues can be used to understand changes in their biological functionality during progression of protein misfolding diseases. We reasoned that it is important to capture proteins in their innate states to preserve the complex cellular milieu without the protein degradation that might occur during extraction and homogenization of organs. In this study, we extended the CPP method to an animal model of disease to probe the dynamic changes in protein structures *in vivo* as AD progresses. This is the first *in vivo* study of structural changes of proteins in progressing AD on a proteome-wide scale in mouse tissue. We identified proteins whose structures were altered in co-expressed protein communities across 7 types of mouse tissue, which helps us understand the role of spatially altered proteins in various biological processes.

## MATERIALS AND METHODS

### Animals/Tissue collection

Female mice (APP^(NL-F)^)^21^ were purchased from RIKEN Brain Science Institute and female C57BL/6 were obtained from The Scripps Research Institute breeding colony. Mice were housed in plastic cages located inside a temperature- and humidity-controlled animal colony and were maintained on a standard cycle (a 12 h day/night cycle). Animal facilities were AAALAC (Association for Assessment and Accreditation of Laboratory Animal Care) approved, and protocols were in accordance with the IACUC (Institutional Animal Care and Use Committee). Mice were sacrificed at 6, 9, 12, and 15 months of age.

### First dimethyl-labeling and tissue collections

Mice were anesthetized by inhalation of 1% isoflurane. Chests of the anesthetized mice were opened by cutting the ribcage. The left heart ventricle was punctured with a perfusion needle and a small cut was made in the right atrium to allow outflow of the perfusion solutions. Blood components were washed away with prewarmed pH 7.4 phosphate-buffered saline (PBS) for 10 min. The mice were perfused with 20 mL of fixation solution (1% CD_2_O) at a flow rate of 2.0 mL/min. Immediately afterward, 40 mL of the solution for the first light-dimethylation reaction (0.3 mM NaBH_3_CN, 1% CD_2_O in pH 7.4 PBS) was added at a flow rate of 2.0 mL/min. Organs were quickly excised and cut into 50 mg of tissue blocks. The tissue blocks were incubated in the same labeling solution (0.3 mM NaBH_3_CN, 1% CD_2_O in pH 7.4 PBS) for 10 min, and then the reaction was quenched by immersing the tissue blocks in 50 mM ammonium bicarbonate (ABC) solution for 5 min.

### Tissue homogenization and protein extraction

Tissue blocks were placed in 100 μL of 20 mM 2-[4-(2-hydroxyethyl)piperazin-1-yl]ethanesulfonic acid (HEPES) pH 7.4 and were homogenized with a pestle until no chunks were visible. The tissue samples were sonicated for 10 cycles (pulse-on 5 sec, pulse-off 3 sec, amplify 30%) and then homogenates were clarified by centrifugation at 8,000 g at 4 °C for 30 min. Protein precipitation was performed by adding 400 μL of 100% methanol, 100 μL of 100% chloroform and 300 μL of water to the sample. After vortexing vigorously, the samples were centrifuged at 8,000 g at 4°C for 30 min. The large aqueous layer was discarded. The samples were washed by adding 800 μL of 100% methanol and vortexing vigorously. After centrifugation at 8,000 g at 4 °C for 30 min, the supernatant was removed. The methanol washing step was repeated 3 times. Methanol was removed and the pellet was air-dried. The pellet was dissolved in 100 μL of 1% sodium deoxycholate (SDC) in 20 mM HEPES pH 7.4. The protein concentration was determined with a BCA protein Assay kit following the instructions from the vendor (23225, Thermo Scientific).

### Proteolysis of labeled proteins with chymotrypsin

Aliquots of tissue samples that contained 200 ug of proteins were adjusted to 80 μL with 1% SDC in 20 mM HEPES pH 7.4. The proteins were reduced with 10 mM TCEP (Tris(2-carboxyethyl)phosphine hydrochloride) and 1% SDC in 20 mM HEPES pH 7.4 at 60°C for 60 min on a shaker. Reduced proteins were alkylated with 20 mM IAA (iodoacetamide) for 30 min at 25°C in the dark. Denatured proteins were digested with chymotrypsin (Promega) at 1:100 (enzyme:substrate(w:w)) at 37°C for 16 hr. Samples were acidified with formic acid to a final concentration of 1%. The sample was centrifuged at 8,000 g at 4°C for 30 min and the supernatant was transferred to a new tube. The sample was centrifuged again at 8,000 g at 4°C for 30 min to collect the clean sample and the supernatant was transferred to a new tube.

### Second dimethyl-labeling and desalting

Pierce C18 spin tips (87784, ThermoFisher) were used for the second dimethyl-labeling step and desalting. A multipipette and a 96-well plate were used to prepare multiple samples in one batch. The C18 tips were activated by aspirating and dispensing 100 μL each of 100% methanol and 100% acetonitrile (ACN). After the C18 tips were washed with 100 μL of 0.1% formic acid, the samples were loaded onto the C18 tips by aspiration. To clean the samples bound to C18 tips, 100 μL of 0.1% formic acid was aspirated and dispensed, and the pH was adjusted by aspirating 20 mM HEPES pH 7.4. Peptides bound to the C18 tips were dimethyl-labeled by aspirating 1% formaldehyde (^13^CD_3_O), 0.3 mM Sodium cyanoborodeuteride (NaBD_3_CN) and the saturated tips were incubated for 15 min at 25°C. The reaction was quenched by aspirating 50 mM ABC and incubating for 10 min at 25°C. After washing C18 tips with 0.1% formic acid, the labeled peptides were eluted with 100 μL of 40% ACN in 0.1% formic acid followed by 100 μL of 60% ACN in 0.1% formic acid. The eluted peptides were lyophilized.

### Strong cation exchange (SCX) fractionation of peptides

SCX fractionation was conducted with commercial spin columns (90008, ThermoFisher Scientific). The pH of the sample was reduced by adding 800 μL of 30% ACN in 0.1% formic acid. The spin column was equilibrated by adding 400 μL of 30% ACN in 0.1% formic acid. It was then centrifuged at 1,000 g for 5 min, and the flow-through solution was discarded. The sample was loaded on the spin column and was centrifuged at 1,000 g for 3 min. Flow-through was stored for LC-MS/MS analysis. The peptides were eluted with consecutive 200 μL aliquots of elution buffer containing 10 mM, 30 mM, 50 mM, 70 mM 100 mM, 150 mM and 300 mM of ammonium acetate. All elution buffer aliquots contained 0.1% formic acid and 30% ACN.

### LC-MS/MS analysis

Samples were loaded onto EvoTips following the manufacturer’s protocol. The samples were run on an Evosep One (Evosep) coupled to a timsTOF Pro (Bruker Daltonics). Samples were separated on BEH 1.7 μm C18 beads (Waters) packed in a 15 cm × 150 μm inner diameter column with an integrated tip (pulled in-house) using the 30 SPD (samples per day) method. Mobile phases A and B were 0.1% formic acid in water and 0.1% formic acid in acetonitrile, respectively. MS data was acquired in PASEF mode with one MS1 survey TIMS-MS and PASEF MS/MS scans acquired per 1.1 s acquisition cycle. Ion accumulation and ramp time in the dual TIMS analyzer was set to 100 ms each and the ion mobility range spanned from 1/K_0_ = 0.6 Vs/cm^2^ to 1.6 Vs/cm^2^. Precursor ions for MS/MS analysis were isolated with a 2 Th window for m/z < 700 and 3 Th for m/z > 700 with a total m/z range from 100 to1700. The collision energy was lowered linearly as a function of increasing mobility starting from 59 eV at 1/K_0_ = 1.6 VS/cm^2^ to 20 eV at 1/K_0_ = 0.6 Vs/cm^2^. Singly charged precursor ions were excluded with a polygon filter, and precursors for MS/MS were picked at an intensity threshold of 2,500, target value of 20,000 and with an active exclusion of 24 s.

### Peptide identification and quantification

Raw files were searched against mouse proteins from Swiss-Prot-Uniprot database (retrieved 03/13/2022, 51,076 entries) containing canonical and isoform sequences, using MSFragger (version 17.1) in the FragPipe pipeline with mass calibration and parameter optimization enabled.^22^ Philosopher was used to filter all peptide-spectrum matches. Quantification analysis was performed with IonQuant. The parameter setting of chymotrypsin allowed for two missed cleavage sites and the minimal required peptide length was set to six amino acids. Dimethyl peptide pairs were identified using variable modifications of light (Δ mass: 32.0564) and heavy labeling (Δ mass: 36.0757) of lysine, oxidation of methionine (Δ mass: 15.9949), fixed modification of heavy dimethylation (Δ mass: 36.0757) on N-terminus and the carbamidomethylation of cysteine (Δ mass 57.0214). Precursor tolerance was set to 50 ppm and fragment tolerance was set to 50 ppm. Isotope error was set to 0/1/2. The minimum number of fragment peaks required to include a PSM (peptide-spectrum match) in modeling was set to two, and the minimum number required to report the match was four. The top 150 most intense peaks were considered, and a minimum of 15 fragment peaks were required to search a spectrum. The data were also searched against a decoy database and protein identifications were accepted at 1% peptide false discovery rate (FDR). All identified peptides were heavy-dimethylated on N-terminus.

### Determination of accessibility of lysine sites

Each peptide with a lysine site should be either light- or heavy-dimethylated, depending on the accessibility of lysine site. The difference in intensity of the peptides labeled in the first isobaric labeling step versus the second yields a relative abundance ratio R.^23^ The R value represents the proportion of the peptide in which a specific lysine site was accessible for dimethylation and is independent of the overall protein amount in the sample.^19^ The relative accessibility of a lysine residue for dimethylation is assessed by the value of accessibility; Accessibility (%) = R/(1+R) × 100.

### k-Nearest Neighbor (kNN) Imputation

Missing values were imputed by kNN machine learning method using the VIM package in R.^24, 25^ The kNN method assumes a relationship between spot volume patterns of groups of proteins. The kNN method impute missing values by selecting spots with spot volume patterns similar to the spot of interest.^26^ A weighted average of values from the k most similar spots is used as an estimate for the missing value. The contribution of each spot is weighted by its similarity determined as the Euclidean distance. The optimum number of k-neighbors must be determined empirically.

### Weighted correlation network analysis (WGCNA)

A weighted protein co-expression network was built using the value of protein abundance from blockwiseModules WGCNA function.^27^ Construction of weighted gene co-expression networks was conducted independently for each of 7 tissue datasets. The soft thresholding powers were determined with the R function pickSoftThreshold.^28^ To pick an appropriate soft-thresholding power for network construction, the value of power was raised to 50. The chosen values were the smallest threshold that resulted in a scale-free R^2^ fit of 0.75 and the networks were created by calculating the component-wise minimum values for topologic overlap. Soft threshold powers varied across seven tissues as follows: 22 for brain, 16 for heart, kidney, 12 for liver, muscle and spleen, and 26 for thymus. BlockwiseModule function was run with the following parameters: TOMType = “signed”, maxBlockSize = 5000, mergeCutHeight = 0.1, verbose = 3. Module eigenprotiens (MEs) were calculated the correlation between the traits of AD (AD vs. NC). Multiple comparisons were accounted for by FDR correction across modules, and the *P*-values for the modules were reported.

### Statistical analysis

Differentially expressed proteins and altered accessibilities (%) between pairs of different age groups (6, 9, 12, and 15 months) in AD or between different pathological conditions (AD and NC) per age were found using Mann-Whitney tests independently. Kruskal-Wallis was used to simultaneously compare the accessibilities among four age groups (6, 9, 12, and 15 months) or among three age groups (6, 9, and 12 months or 9, 12, and 15 months) in AD. These comparisons were tested with Kruskal-Wallis followed by Bonferroni’s comparison *post hoc* test independently. The criterion for significance was a *P*-value less than 0.05.

### Enrichment of Gene ontology (GO) and protein-protein interaction analysis

GO analysis was performed with ClueGO (a plug-in in Cytoscape) to identify the significant biological functions of the proteins in the WGCNA module.^29^ Protein-protein networks were detected using Metascape and the following databases: STRING, BioGrid, InWeb_IM, OmniPath.^30–32^ The resultant network contained the subset of proteins that form physical interactions with at least one other list member, the confidence cutoff of physical interaction was set to medium (0.5) or strong (0.7). Visualization of the protein-protein interaction network was performed on the Cytoscape combining STRING.

### Complex modeling with AlphaFold2-multimer

We used AlphaFold2-multimer to predict the protein-protein interaction motif of each complex. AlphaFold2-multimer modeling was performed with ColabFold.^33^ Input multiple sequence alignment (MSA) features were generated by local ColabFold using the “MMseqs2 (Uniref[+[Environmental)” MSA mode. By default, the constructed MSAs contain both unpaired (per-chain) and paired sequences. AlphaFold2-multimer was run with one or several options from the following list: model type = alphafold2_multimer v3, num recycles = 3, recycle early stop tolerance = 0.5, max msa = auto, num seeds = 1. The models were ranked by confidence score, and rank 1 was selected as the most accurate model. The distance between two lysine residues was calculated using PyMOL2 version 2.5 (Schrödinger, LLC).

## Results

### Deep profiling of dimethyl-labeled peptides in mouse tissues

To probe the structural changes of the proteome in tissues in progressing AD, we developed a method for *in vivo* dimethyl labeling of proteins. Formaldehyde solution was diffused through blood vessels and into tissues by perfusion through the heart. All solvent exposed lysine residues were labeled with formaldehyde to form a Schiff’s base. This initial step was immediately followed by perfusing a cyanoborohydride solution into the mouse to convert the Schiff’s base to a dimethyl label. The process is fast enough to capture protein structures in a nearly innate state. Exposed lysine sites on the surface of proteins were light-dimethylated [(CHD_2_)_2_] and then 7 organs were harvested. After homogenization of each tissue and lysis of the cells, proteins were denatured and proteolyzed with chymotrypsin followed by labeling of the newly exposed lysine sites with heavy-dimethyl [(^13^CD_3_)_2_] tags (Figure 1A). The accessibility of each lysine site was determined from the ratio of the intensity of light-labeled peptide vs. the sum of intensities of the light- and heavy-labeled peptides. We systematically investigated the structural changes in the proteomes of 7 tissues in an AD mouse model ranging in age from 6 to 15 months, as well as in normal control (NC) mice to exclude the effects of aging.

**Figure 1.**
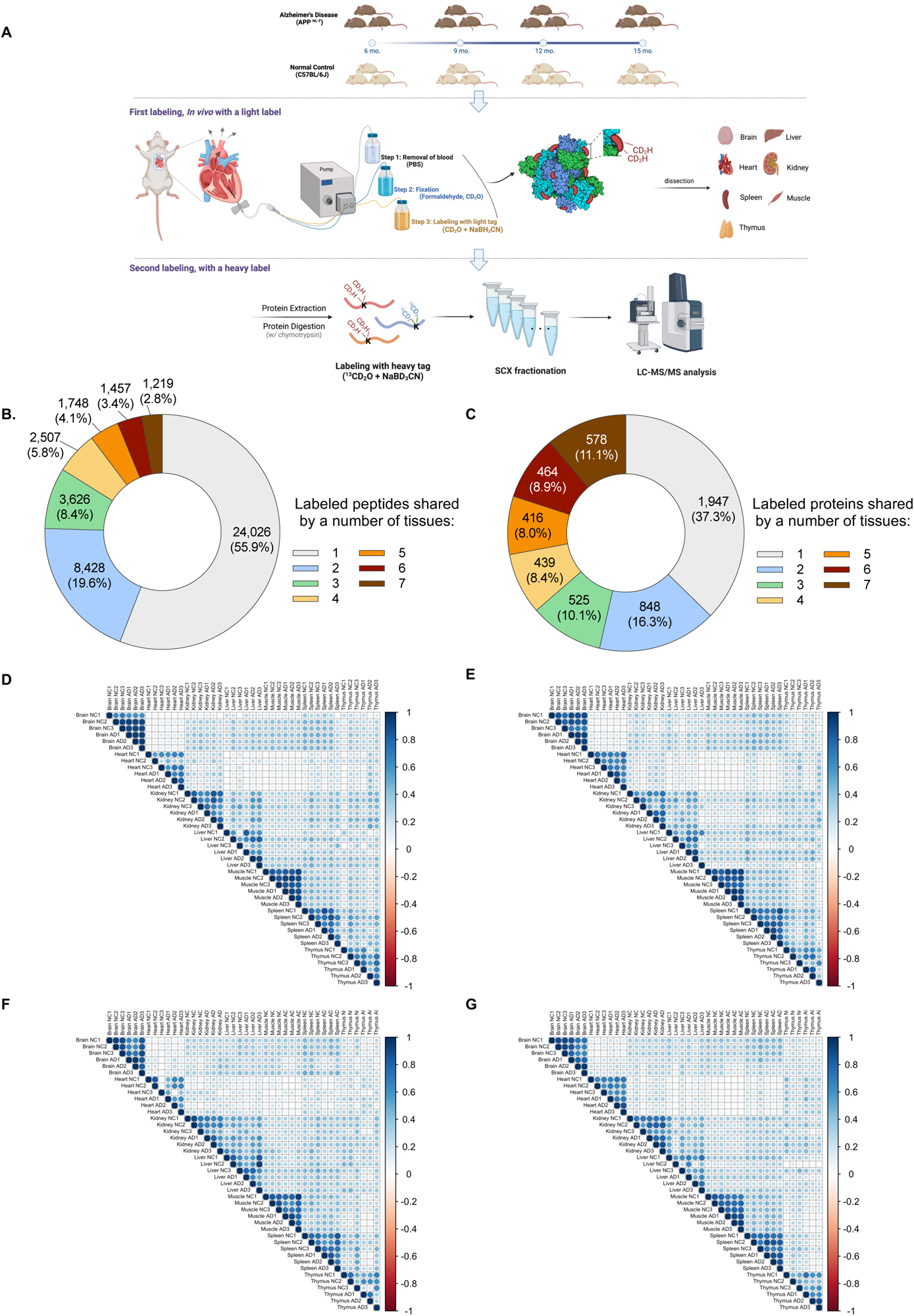
Strategy for identifying dimethyl-labeled peptides. **A.** Three mice per each age group (ranging 6 months to 15 months) were used for AD (APP^NL-F^) and NC (C57BL6/J). The first step of the CPP workflow consists of three sub-steps that were conducted via perfusion: i) blood was washed by PBS, ii) tissue was fixed by formaldehyde, and iii) exposed lysine sites of the native proteins were labeled with light-dimethylation ([CD_2_H]_2_). Proteins from each of the seven organs were extracted and digested separately with chymotrypsin, after which the newly exposed lysine sites were labeled with heavy dimethylation ([C^13^D_3_]_2_). **B.** More than half of the total labeled peptides were tissue-specific. Less than 3% of a total of labeled peptides were peptides common to all seven tissues. **C.** The proportions of labeled proteins were determined by assigning proteins to the labeled peptides. Unlabeled proteins were not counted. The largest portion of labeled proteins was tissue-specific proteins, and the portion of proteins common to all 7 tissues was the third largest portion. **D-G** Biological triplicates were correlated across 7 tissues at 6 months (D), 9 months (E), 12 months (F) and 15 months (G).

A total of 43,014 dimethyl-labeled peptides that mapped to 5,217 proteins across all tissues were identified at a peptide false discovery rate (FDR) of < 1% (Figure 1B and 1C). Among the labeled peptides, 1,219 labeled peptides that mapped to 498 proteins were identified in all 7 tissues, whereas 24,026 labeled peptides that mapped to 4,952 proteins were tissue-specific (Figure 1B). Likewise, the highest proportion of labeled proteins were tissue-specific (37.3% (n = 1,947)), whereas labeled proteins that were identified in all 7 tissues comprised 11.1% (n = 578) of all labeled proteins (Figure 1C).

We next sought to examine the reproducibility of the *in vivo* CPP method for each mouse tissue. We evaluated the correlations of the accessibility between biological replicates for each age, then averaged these correlations. We found the highest R values for NC (0.842) and AD (0.782) in brain, whereas thymus showed the lowest correlation, 0.549 for NC and 0.505 for AD (Figure 1D-G, Table S1). Correlations averaged across all ages in all 7 tissues showed a correlation in accessibility, with R values of 0.668 and 0.649 for NC and AD, respectively. Because we observed strong correlations of the biological replicates regardless of anatomical source or pathological conditions of the tissue, we concluded that perfusion-based dimethyl-labeling was reliable. Also, we noticed that correlations of accessibility were higher within the same tissues, even in the presence of different pathological conditions, compared to those observed between different tissues at the same age.

### Formaldehyde and cyanoborohydride diffused evenly in first labeling reaction

In CPP, dimethyl labeling relies on the reductive amination of the lysine residue.^19, 34^ Although the formation of the Schiff’s base is fast, the perfusion process requires about 30 minutes for completion. We sought to examine whether the distribution of reagent via blood vessels was even and how efficiently proteins were dimethylated across seven organs. Efficiency of perfusion was assessed based on the proportion of K-containing peptides that were not light-dimethylated on lysine. Of a total of 23,039 peptides detected in the brain, 39.0% of peptides (n = 8,974) harbored lysine sites, and 8,758 of the 8,974 peptides were labeled with light- or heavy-dimethyl modifications (or both) (Figure 2A). Of a total of 4,295 detected proteins in brain, 2,565 (59.7%) proteins were modified with dimethyl. Among the 8,974 lysine-containing peptides, 8,368 were light-dimethyl labeled through perfusion, indicating a high labeling efficiency (93.2%), while 606 peptides (6.8%) were not labeled with light-dimethyl tags (Figure 2B). Among the 606 peptides that were not light-dimethylated, 390 were solely heavy-dimethylated and 216 were not labeled with either light- or heavy-dimethyl tags. Heart showed the highest efficiency of light-dimethylation, 10,745 peptides (97.5%) of 11,018 K-containing peptides. We presume that the high light-labeling efficiency of heart is due to the extended time that reagent remained in heart before being circulated to other organs. Labeling efficiency was high and consistent across all organs, ranging from 91.5% in muscle to 97.5% in heart, with a standard deviation for all organs of 2.3%. The average of the first labeling efficiency in all 7 tissues was 94.8%. Thus, we concluded that the first labeling via perfusion was efficient, and the reagents evenly diffused across all tissues.

**Figure 2.**
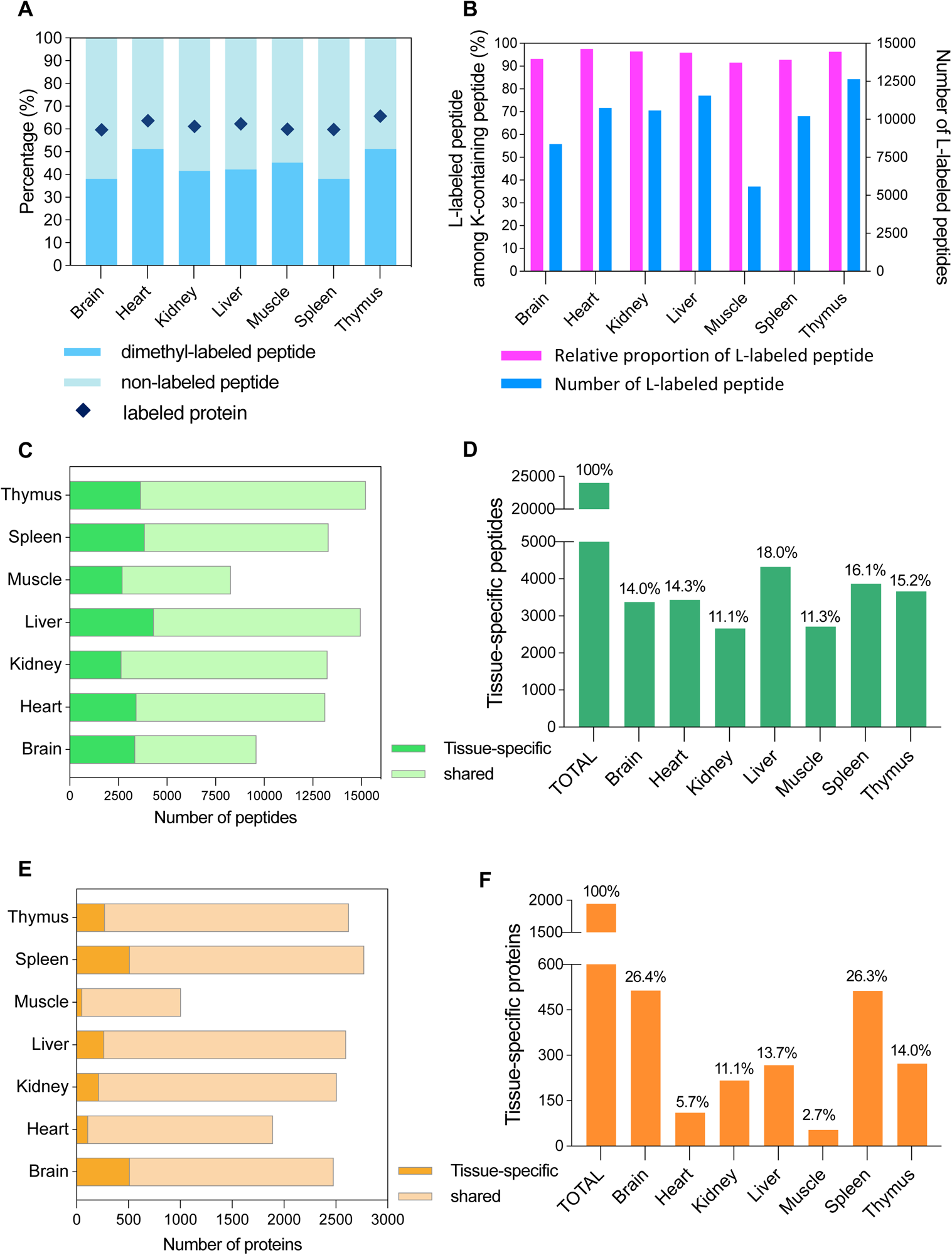
Labeling efficiency of the 1^st^ labeling via perfusion. All the labeled peptides (light, heavy or both light and heavy) were counted as identified peptides for each tissue sample. The proportions of identified peptides that were labeled ranged from 38.01% (n = 8,758) in brain to 51.20% (n = 10,779) in heart (blue). Diamonds indicate the relative proportion of proteins that were labeled. The proportions of lysine-containing peptides that were light-labeled were determined for each tissue. The proportions ranged from 91.5% (n = 5,583) in muscle to 97.5% (n = 10,745) in heart (pink). All tissue samples show a labeling efficiency of more than 90% for labeling via the perfusion method, while the number of light-labeled peptides were variable (blue). **C, E.** The tissue-specific labeled peptides (C)/proteins (E) (green/orange) and shared peptides/proteins (pale green/pale orange) were determined for each tissue sample. **D, F.** These bar graphs represent the contribution of each tissue to the total number of tissue-specific labeled peptides (D)/proteins (F).

We then plotted the dimethylation pattern in each tissue. The liver had the second highest number of identified labeled peptides (n = 14,962) but the highest number of tissue-specific labeled peptides (n = 4,321), accounting for 18% of all 24,026 tissue-specific labeled peptides (Figure 2C and 2D). The kidney had the lowest number (11.1%, n = 2,664) of labeled tissue-specific peptides but ranked third lowest in the number of labeled tissue-specific proteins (11.1%, n = 216) (Figure 2F). Muscle-specific proteins represented a small proportion (2.7%, n = 53) of tissue-specific proteins out of a total of 1,947 (Figure 2E and 2F). However, the presence of highly abundant proteins in muscle may have affected the identification of lower abundance proteins. The top 100 muscle proteins accounted for 78.6% of all identified muscle proteins, while the top 100 proteins in the kidney and heart accounted for 37.8% and 57.5% of the identified proteins, respectively (Figure S1A). Myosin regulatory light chain 11 (Mylpf), the most abundant protein in muscle, contributed 11.3% of the total muscle protein mass, and the top 12 proteins in muscle represented more than half of the mass of the top 100 proteins (Figure S1B). In conclusion, the labeling data provides tissue-specific protein information, which is valuable for understanding the physiological changes associated with disease in each tissue.

### Variability of the conformational changes among 7 tissues

We used labeled peptides that were detected in all seven tissues to quantitatively measure the structural differences in proteins. To minimize the effect of tissue-biased accessibility, we used quantile-normalized values for comparison across tissues (Figure S2). We sought to identify patterns of changes in accessibility that occur specifically for AD in proteins from 6mo to 15mo. To achieve this, we tested for differences over time between NC and AD using spline model from 6 to 15 months for each labeled peptide (Figure 3A). Brain tissue was the most structurally affected by AD, with 686 peptides in AD brain showing significantly different patterns of accessibility compared to NC from 6 to 15 months (Figure 3B). In heart, kidney, and thymus, fewer than 25 peptides showed significantly different patterns of change in AD relative to NC. There were no labeled peptides that exhibited significant different patterns in accessibility across all organs from 6 months to 15 months. However, a total of 10 labeled peptides consistently showed significantly different patterns in accessibility across four different organs each in the AD model. To examine the conformational changes of proteins specifically impacted by AD, we corrected for the confounding effect of age by dividing the individual accessibility of AD by the accessibility of its corresponding sequence of NC, resulting in a metric referred to as “fold-change” in this study. We evaluated the variability in structural changes depending on tissues relative to the brain using 10 peptides that showed distinct patterns of change in accessibility in AD from those in NC in four tissues (Figure 3C-F, Table S2). At the early stage of AD (6 months), the biggest conformational discrepancy due to AD was observed between the liver and brain. The lysine site of TAKGLF (Eno1) was not only more accessible in AD liver than in AD brain, but also more accessible in AD liver than in NC liver. As AD progressed, AD-specific structural changes in muscle and spleen were greater than those in brain when the fold-change between each tissue and the brain was compared. (Figure 3G-H). A functional enrichment analysis of the 10 proteins retrieved KEGG pathways associated with metabolism, glycolysis, and the TCA cycle (Figure 3I). For only 5 (51-67 amino acid (AA) of Atp5f1d, 469-479 AA of Dpysl2, 48-57 AA of Eno1, 1472-1483 AA of Flna, and 277-284 AA of Ppp2cb) out of 1219 peptides detected commonly in the 7 tissues, the accessibility patterns in all 7 tissues during AD progression were not significantly different from those during normal aging, and no significant difference in accessibility was observed across the 7 tissues under each condition (4-ages, disease). This indicates that the regions corresponding to these 5 peptides were not affected by AD in all 7 tissues. As expected, this investigation confirms that the effects of AD are most significantly observed in brain tissue, but by quantifying conformational changes of proteins across all tissues, we found tissue-specific variations associated with AD.

**Figure 3.**
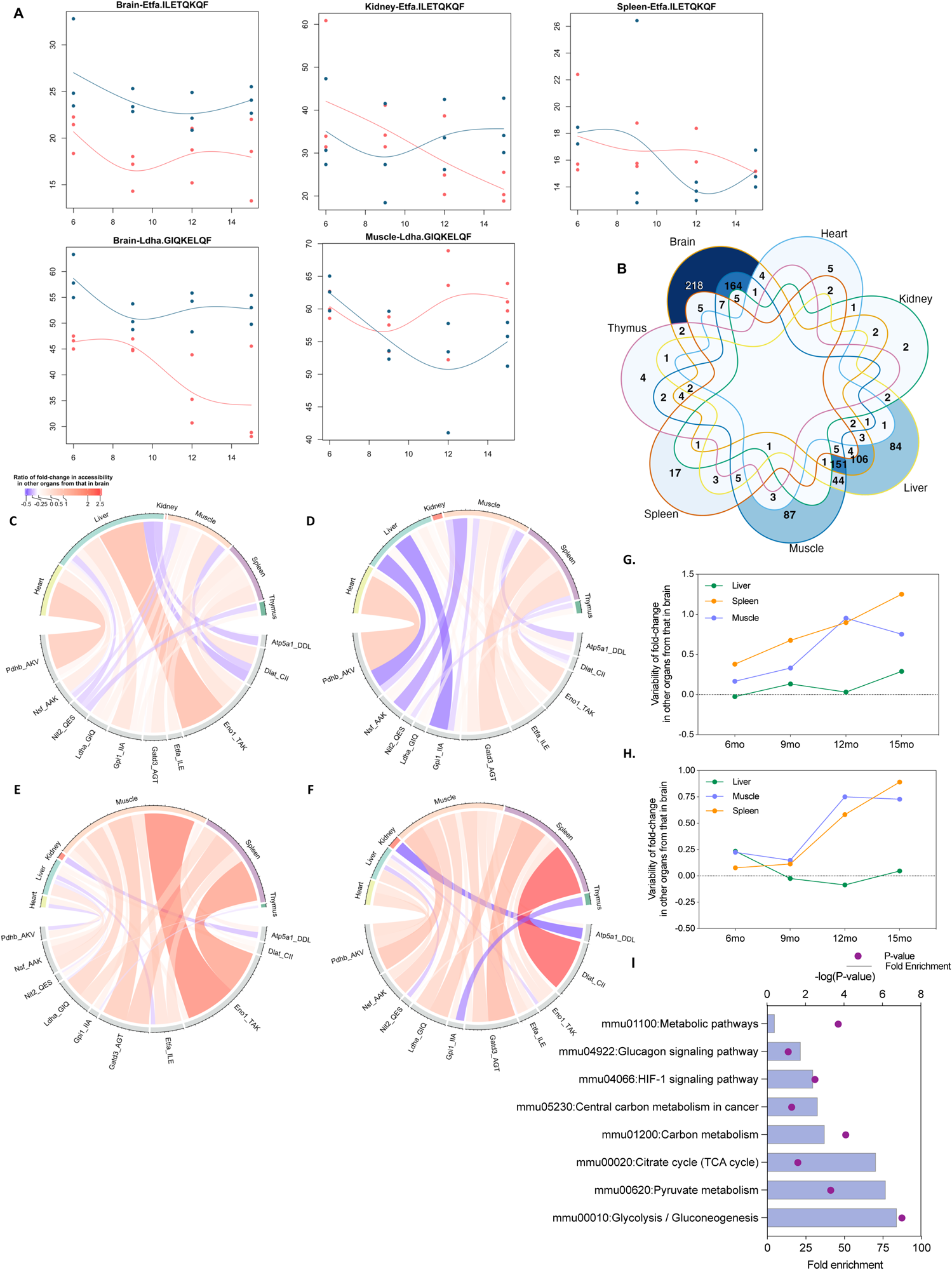
Variability of the conformational changes depending on the tissues. Two representative peptides (ILETQKQF and GIQKELQF) were shown representatively. The normalized values were utilized to fit spline models. The accessibility of each peptide for both NC (blue) and AD (red) exhibited significantly distinct patterns from 6 to 15 months (P-value < 0.05). Venn diagram shows the number of peptides exhibiting significant differences in the trend of accessibility changes between NC and AD. There were no peptides from AD that showed a significant difference in accessibility changes compared to NC in all 7 tissues during the period from 6 to 15 months. The value of zero was not indicated. **C-H** During AD progression, 10 common peptides exhibited distinct patterns in accessibility between NC and AD in different four tissues. The variabilities for the structural changes in each tissue were calculated based on the value of brain using the formula: (fold-change of other tissue - fold-change of brain) / fold-change of brain at 6 mo (C), 9 mo (D), 12 mo (E), and 15 mo (F). Only the first three amino acids were shown. AGTAEAIKAL of Gatd3 (G) and GIQKELQF of Ldha (H) showed a difference in the magnitude of accessibility change in muscle and spleen compared to that in the brain as AD progressed. **I.** Enriched KEGG pathways with 10 proteins.

### Conformations of proteins in the brain are changing as AD progresses

Proteomic investigations into AD pathology have primarily relied on the analysis of differential protein expression. Bing et al. profiled the differentially expressed proteins and identified the protein networks that are affected during AD progression.^35^ Co-expressed proteins and altered protein expression in human brain tissue of asymptomatic and symptomatic AD patients were reported by Nicholas et al.^36^ Savas et al. measured protein expression in several mouse models of AD using quantitative mass spectrometry.^37^ Despite extensive proteomic studies on AD and some footprinting studies on AD related proteins there have been no comprehensive *in vivo* studies of protein structures as AD progresses.^19, 38–40^

To uncover the changes in the 3D structure of brain proteins, we focused on 780 proteins, which are known to be expressed in the brain based on the Human Protein Atlas database (https://www.proteinatlas.org) and were also found to be dimethyl-labeled in our brain dataset. To identify lysine sites that change significantly in accessibility during the progression of AD and which also differ from NC, we used statistical methods to test the accessibility of 3,456 peptides corresponding to 780 proteins. First, a Kruskal-Wallis test was used to simultaneously compare lysine site accessibility among the four age groups (6, 9, 12, and 15 months) of AD (P-value ≤ 0.05), and then Mann-Whitney was used to compare lysine site accessibility between AD and NC per age (P-value ≤ 0.05). Of the 3,456 peptides that were tested, 83 peptides corresponding to 62 proteins showed a significant difference in accessibility in all tests. These results suggest that the accessibility of these peptides changed consistently during the progression of Alzheimer’s disease, and that they simultaneously diverged from the NC, thus indicating that the structural changes observed in the AD samples were not induced by aging (Figure 4A, 4B). Four pairs of peptides that shared lysine sites provided validation for the measured changes: APVISAEKAY and APVISAEKAYHEQL for Tuba1, HPEQLITGKEDAANNY and ITGKEDAANNY for Tuba1, QVVLVEPKTAW and QYQVVLVEPKTAW for Cnp, and RYLSEVASGENKQTTVSNSQQAY and SEVASGENKQTTVSNSQQAY for Ywhab. Many lysine sites showed a tendency to become inaccessible as aging progressed in both the NC and AD groups. However, the lysine sites in the AD group were found to be more inaccessible than those in the NC group. This suggests that the conformation of proteins in AD may be altered by the physiological changes associated with the disease. For example, 2’,3’-cyclic-nucleotide 3’-phosphodiesterase (Cnp), which is associated with neuronal cells and glial cells,^41^ was found to be enriched in the brain. When compared to the average expression levels in other tissues, the increase in expression levels of Cnp in the brain ranged from 13.4- to 23.9-fold in the NC and from 17.0- to 27.7-fold in the AD across four age groups (Figure 4C). The accessibility of 2 peptides of Cnp showed a decrease pattern in progressing AD but remained unchanged in the aging NC (Figure 4D and 4E). Of the 62 proteins that showed a significant difference in accessibility, 60 proteins (excluding two brain-specific proteins) exhibited a range of enrichment factor from 0.13-fold (Eef2) to 1,232-fold (Tuba1b) when compared to their expression in other tissues (Figure 4F). Specifically, on average, 51 of these proteins were expressed at higher levels in the brain than in other tissues, while 9 proteins were expressed at lower levels in the brain than in other tissues. Expression of a total 60 proteins was also enriched 21.7-fold in NC and 20-fold in AD on average, but no tendency in expression was observed in either aging or AD status.

**Figure 4.**
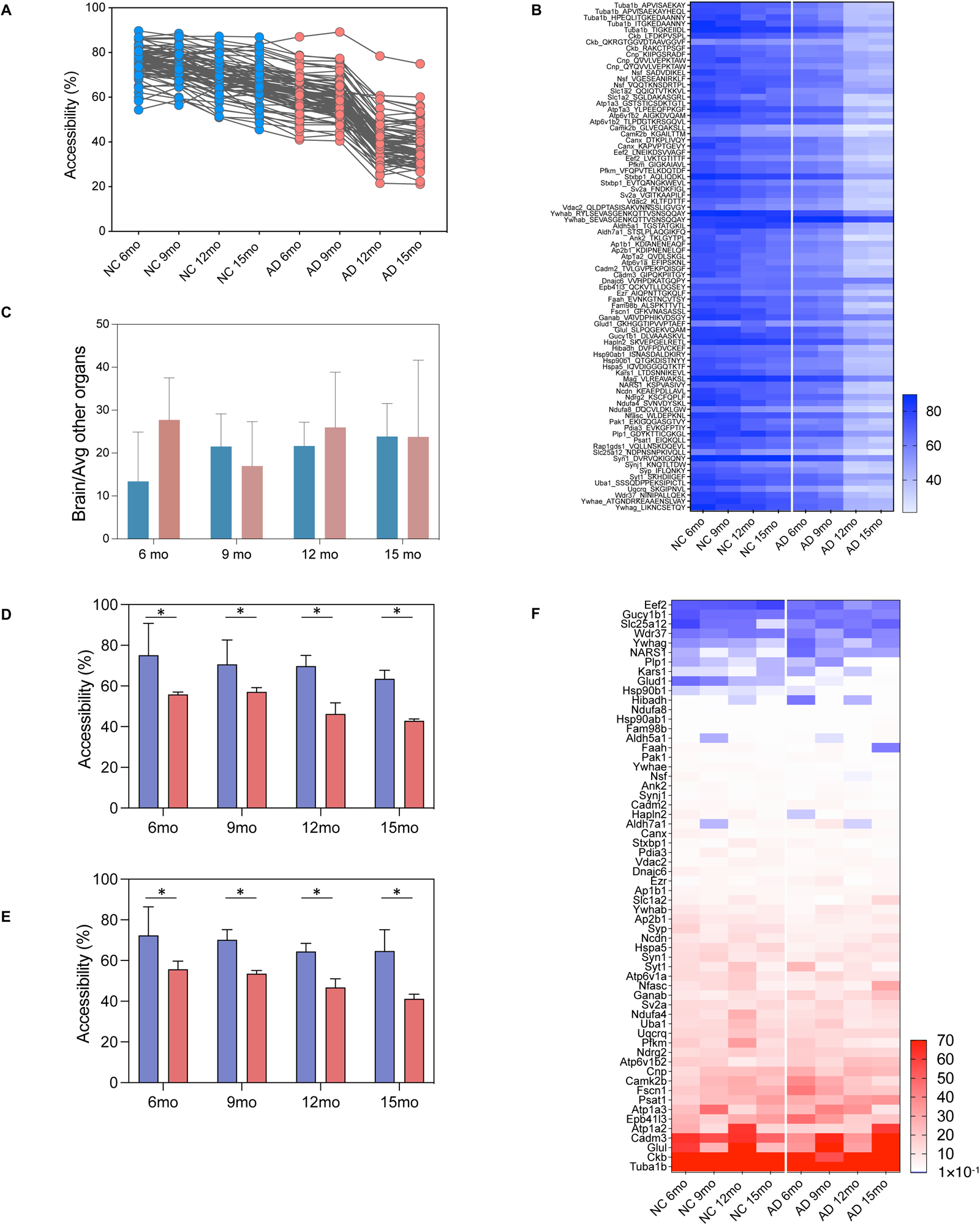
Changes in the structures of proteins that are known to be associated with brain. **A, B.** Accessibility of 83 labeled peptides that mapped to 62 proteins were significantly changed. The accessibility values in AD groups decreased more steeply than those in NC groups (A). The proteins and peptides corresponding to each change in the accessibility are indicated (B). **C, D, E.** Cnp was more highly expressed in brain than in the other tissues (C). Two of the three peptides of Cnp share one lysine site, and variations in accessibility for these two peptides are represented. KIIPGSRADF (D) is located at 87-96 amino acid of Cnp and QYQVVLVEPKTAW (E) is located at 141-153 amino acid of Cnp. While they exhibited a decreasing trend in AD, the accessibility values were lower in AD compared to NC, and the trend in AD was steeper than in NC. Blue indicates NC groups, pink indicates AD groups. Asterisk (*) denotes significance. **F.** Expression of 60 brain proteins were compared to averaged expression in six other tissues (F). The minimum enrichment factor was 0.13 for Eef2 in NC at 15 mo and the maximum enrichment factor was 1,232 for Tuba1b in NC at 12 mo. Expression of 6 proteins (Eef2, Gucy1b1, NARS1, Slc25a12, Wdr37, Ywhag) was lower in brain than the expression of the corresponding proteins in other tissues at all ages in NC and AD. The bar indicates the enrichment factor, with red indicating an enrichment factor more than 1, and blue indicating an enrichment factor less than 1. Proteins enriched more than 70-fold are marked in red.

These results suggest that the structural information obtained through the accessibility of lysine sites can complement protein expression data by revealing changes that are not fully characterized by protein expression levels alone. This method provides an approach to collect both protein expression and structural information to gain a more comprehensive understanding of the changes that occur in a proteome during the progression of a disease.

### Differential structural changes of tightly regulated proteins

We hypothesized that the changes in protein expression could accompany structural changes to proteins, which may suggest alterations to their physiological function. To evaluate our hypothesis, we initially analyzed the abundance of proteins in tightly co-regulated protein networks by modularizing them into protein communities using a WGCNA algorithm.^28^ WGCNA was applied to the dataset of each tissue; 4,295 proteins in brain, 3,074 proteins in heart, 4,186 proteins in kidney, 4,298 proteins in liver, 1,721 proteins in muscle, 4,800 proteins in spleen, 4,159 proteins in thymus were used to build protein co-expression networks. No outliers were detected after all samples were hierarchically clustered using average distance and Pearson’s method. For brain, the lowest soft threshold power was 22, with an R^2^ of more than 0.75. This network consisted of 17 modules of proteins related by their co-expression across control and disease tissues based on the TOM-based dissimilarity, after merging the modules with dissimilarity (Figure S3A and S3B). The WGCNA analysis also divided the protein data sets into 14, 17, 13, 10, 12 and 10 modules for heart, kidney, liver, muscle, spleen and thymus, respectively (Figure S4A-G).

We evaluated the correlation of the co-expressed proteins with Alzheimer’s disease (AD) by comparing the co-expression of proteins in AD and NC within each module, irrespective of age. Correlation of coefficient R > 0.4 and *P*-value < 0.05 were set as the criteria for significant correlation for the AD trait. We found that only a limited number of protein communities were significantly correlated with the AD trait, with one module (M3) in the brain showing a correlation of 0.41, two modules (M11 and M14) in the kidney showing correlations of 0.43 and 0.44, two modules (M2 and M5) in the muscle showing correlations of 0.44 and 0.46, and one module (M4) in the spleen showing a correlation of 0.44. No modules in other tissues showed a significant correlation with the AD trait. (Figure 5A). We also assessed whether the direction and strength of the association between each module and the AD trait remained consistent (positive or negative) following the subdivision of the samples into four age groups (Figure S4A-G). Strong correlations were observed at a certain age in a few modules, while in most modules, the direction of correlation was inconsistent across the four age groups. For example, when samples from all age groups were included, a high correlation was observed for module 3 (M3) of brain (R = 0.41) (Figure 5A and 5B). Following that, the samples were divided by age. M3 proteins from the 9-month samples were found to be highly negatively correlated with a value of R = 0.82. At 12 and 15 months, samples were also negatively correlated, with R values of 0.42 and 0.72, respectively. In contrast, M3 proteins from the 6-month samples showed a positive correlation with a value of R = 0.4 (Figure S4). The direction of association between the protein abundance-based module and the AD trait was found to fluctuate during the progression of AD. Therefore, we focused on the proteins in modules displaying significant correlations with the AD trait across all samples to investigate the conformational changes of the co-regulated proteins.

**Figure 5.**
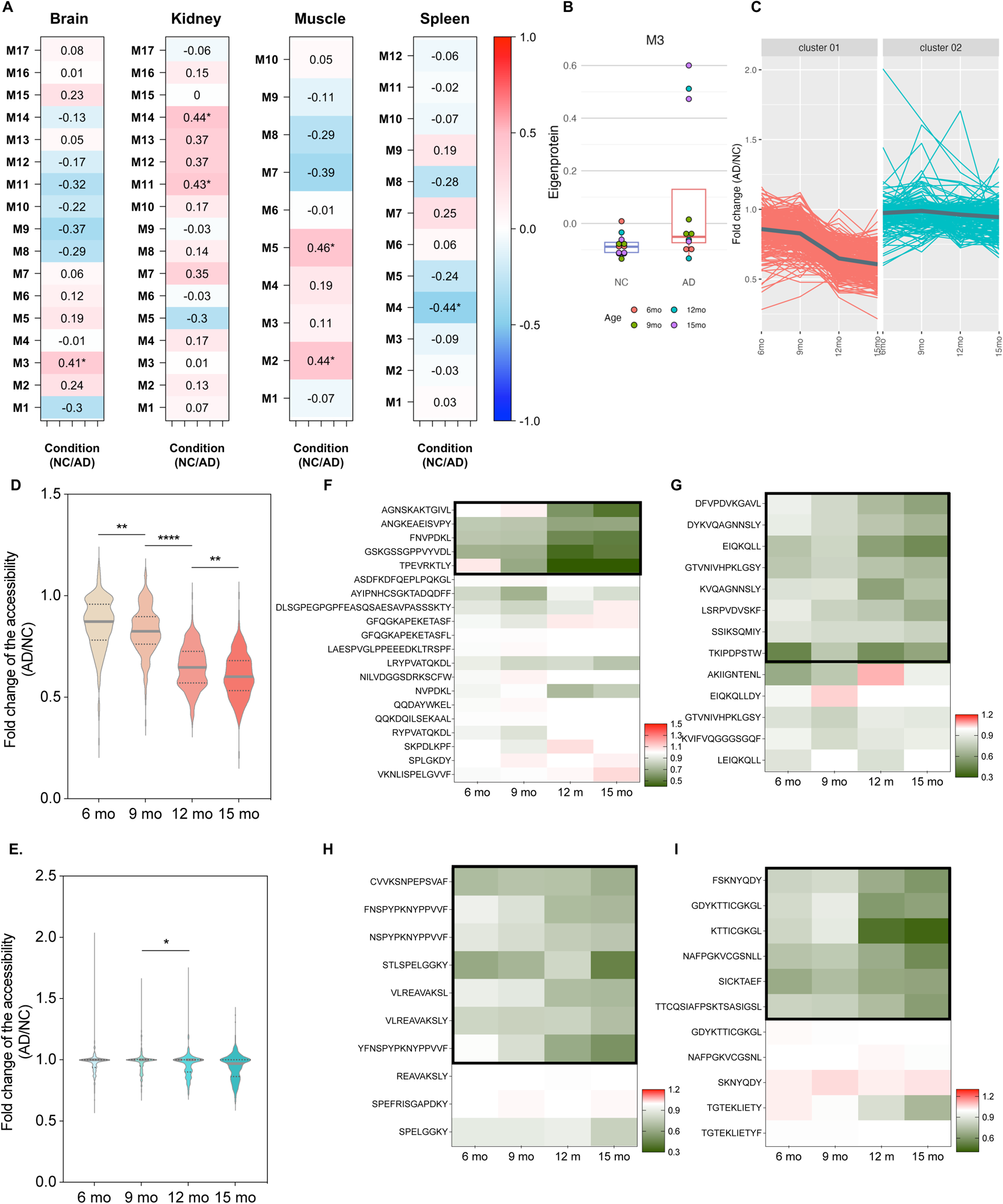
The structural changes of the co-expressed proteins by WGCNA. **A.** Four of the seven tissues (brain, kidney, muscle and spleen) showed significant correlated modules (R^2^ > 0.4, P-value <0.05). **B.** In module 3 (M3) of brain, the eigenprotein level between AD and NC was assessed, with dot colors indicating the age of the mice. **C-E.** The labeled peptides of M3 proteins were clustered based on the fold-change of the accessibility (C). Distribution of the fold-change of the accessibility of cluster 1 (D) and cluster 2 (E). Of 481 labeled peptides that mapped onto 174 proteins in M3, fold-change of accessibility of 268 peptides showed a consistent decrease in progressing AD in cluster 1 and fold-change of accessibility of 213 peptides in cluster 2 did not significantly change in progressing AD. *p < 0.05, **p < 0.005 and ****p < 0.0001 **F-I.** The fold changes of the accessibility for Map1a (F), Psat (G), Mag (H), and Plp1 (I) are presented. The peptides in the bold box were included in cluster 1. Peptides in the bold box clearly decreased.

We investigated the conformational changes of the co-regulated proteins from the 4 types of tissues (brain, kidney, muscle, and spleen) that showed the significantly correlated modules based on WGCNA. To examine whether the conformational changes of proteins were specifically impacted by AD, the labeled peptides were clustered based on the fold-change of the accessibility during the progression of AD using the K-means clustering algorithm. The number of clusters was determined via an optimization algorithm (Figure S5). For the brain, the 481 labeled peptides that were mapped onto 174 proteins constituting M3 were clustered into two clusters (Figure 5C). Accessibility of the lysine sites of 268 peptides (123 proteins) in cluster 1 showed a steadily decreasing pattern, with a slight decrease during early AD development (6-9 months), a dramatic decrease from 9 to 12 months, and a slight decrease again during the late stage of AD development from 12 to 15 months (Figure 5D). This suggests that beginning at 9 months, the lysine sites included in cluster 1 were significantly sterically inaccessible due to AD. On the other hand, lysine sites of 213 peptides (113 proteins) in cluster 2 showed relatively stable accessibility during AD development (Figure 5E). Of 174 proteins in M3, 62 have 2 or more peptides included in both cluster 1 and cluster 2, and all labeled peptides of each of 61 and 51 proteins were exclusively included in cluster 1 and cluster 2, respectively (Figure S6). Twenty peptides that mapped to Map1a were identified most often in M3 proteins (Figure S7), and 5 of 20 peptides were hidden as AD progressed, but 15 lysine sites were spatially stable (Figure 5F). Subsequently, 19- and 14-labeled peptides were mapped to Pkg1 and Mdh1, respectively, and the peptides included in cluster 1 exhibited a consistently decreasing pattern of accessibility. The lysine sites of the peptides of Psat1, Mag, and Plp1, which are highly expressed in neuronal cells and were included in 62 brain proteins that constituted M3, became inaccessible during progression of AD (Figure 5G-I). The datasets from kidney, muscle, and spleen were processed separately to reveal the AD-induced spatial changes of proteins (Figure S8). Collectively, our findings suggest that the steric changes of proteins occur concurrently with changes in co-expression of proteins as AD progresses.

### Biological functions of protein communities whose conformational changes precede expression changes

We sought to uncover how the network undergoing conformational changes was related to biological functions, particularly those involved in neurodegenerative diseases. To achieve this, we examined the interactions of these proteins and their functional implications. In this analysis, we used proteins that had shown altered patterns of accessibility fold-change via K-means clustering. For instance, the proteins that were included in cluster 1 were used for the brain dataset.

We investigated how changes in the structure and expression of proteins could affect the progression of Alzheimer’s disease by examining the pathways and biological functions they are involved in. We performed the analysis for the enrichment network on Metascape (metascape.org), considering the inter-term similarity and intra-term redundancy in the enriched terms.^30^ This analysis represented an enriched term as a node connecting other nodes considering Kappa similarities. A total of 113 significantly enriched terms were grouped into 20 clusters based on their similarities and redundancies in the brain dataset (Figure S9 and Table S3). Fifteen proteins (Gnas, Mdh1, Ogdh, Pgk1, Ppp1cb, Slc1a3, Sod2, Taldo1, Sdha, Oxct1, Ndufa8, Epm2aip1, Aldh1l1, Etfa, and Ugp2) were enriched in “generation of precursor metabolites and energy” (node-107) with the most significantly enriched having a *P*-value of 1.78*10^-9^. “Energy derivation by oxidation of organic compounds” (node-1) was also enriched significantly, but was similar to node 107 with kappa score of more than 0.3. It has been demonstrated that an abnormality of carbon and energy metabolism occurs in neurodegenerative disease since neurons require large amounts of energy to maintain their normal activity, and metabolic decline of the brain contributes to cognitive impairment.^42, 43^ Interestingly, we discovered that structural changes preceded the expression changes in both protein communities “generation of precursor metabolites and energy” and “carbon metabolism” (node-96) (Figure 6A-B, 6D-E and Figure S10). Protein expressions in node-107 and node-96 increased in AD but remained stable in NC during aging. Expression fold-change increased significantly at 15 months, whereas accessibility fold-change decreased significantly from 12 months. We also noted that “metal ion homeostasis” (node-108) was significantly enriched with 12 proteins (Ank1, Calb1, Calb2, Gnas, Itpr1, Prkcb, Slc12a4, Slc1a3, Sod2, Vapb, Fis1, Immt). The homeostasis of metal ions is also known to be essential to maintain the normal function of brain, and abnormally elevated iron in brain is recognized to induce cell death and be a cause of several neurodegenerative diseases including AD.^44, 45^ Zinc also has an essential role in protein binding for enzymatic activity or to modulate synaptic transmission, and abnormal levels of zinc have been reported to be implicated in AD.^46, 47^ The enriched proteins in metal ion homeostasis showed an increased but insignificant pattern of expression, while a significantly decreased pattern in accessibility was observed from 9 months (Figure 6C, 6F and Figure S10).

**Figure 6.**
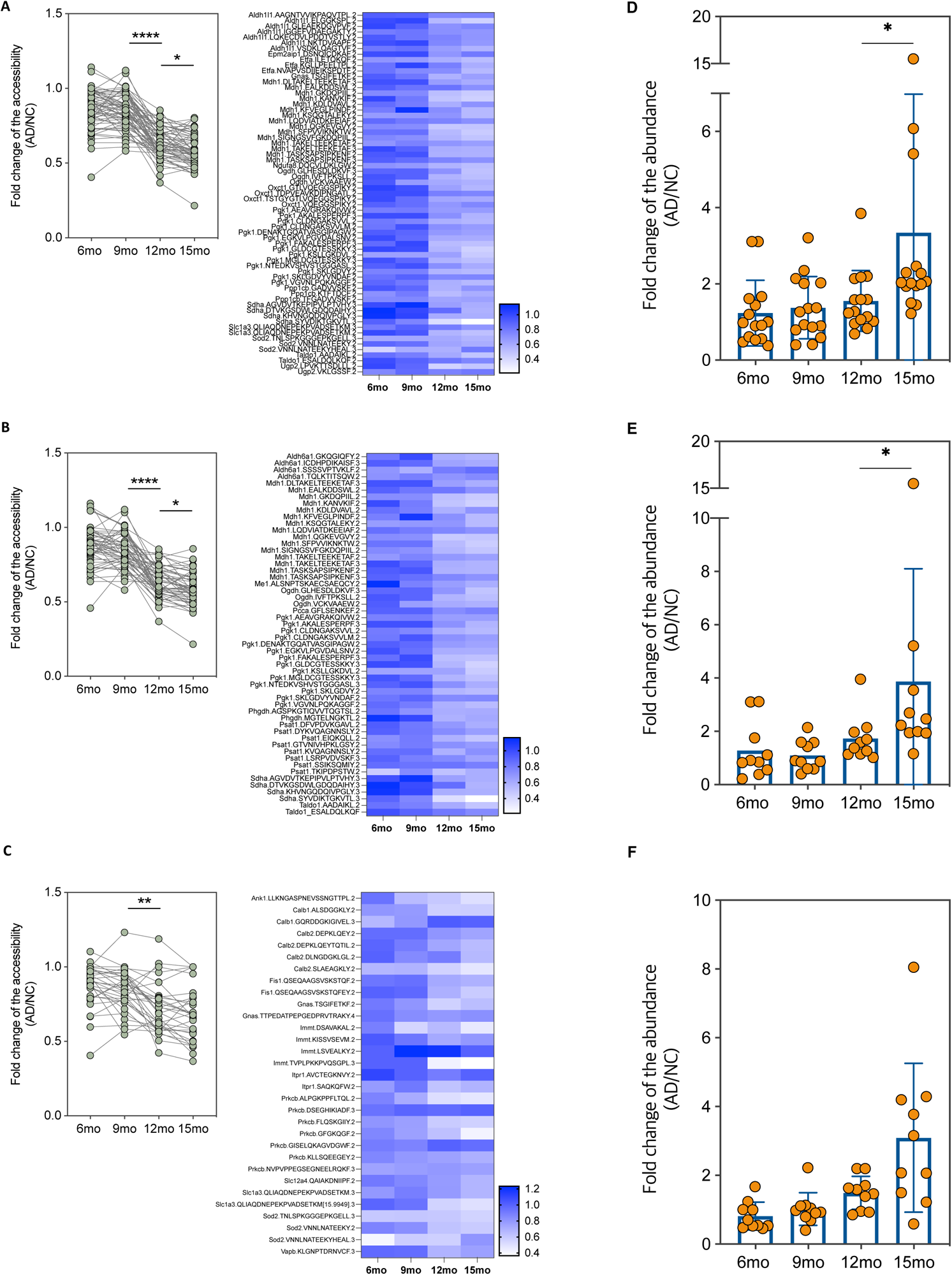
The biological functions affected by structural changes of composed proteins. **A-C.** Structural changes of proteins enriched in generation of precursor metabolite and energy (A), carbon metabolism (B), and metal ion homeostasis (C) are shown. The heatmap (right) showed variations in the fold-change of the accessibility based on peptide sequence. The scatter plots (left) were plotted irrespective of peptide sequences. **D-F.** Expression change of proteins enriched in generation of precursor metabolite and energy (D), carbon metabolism (E), and metal ion homeostasis (F). The fold-changes of the expression level are presented. *p < 0.05, **p < 0.005 and ****p < 0.0001

Next, we used the STRING database to identify 45 out of 123 proteins included in cluster 1 that interacted directly with each other through 30 edges. Of these, 39 proteins were associated with either brain-related terms or the enriched terms in which proteins showed conformational changes preceding the expression change (node-97, node-107, and node-108) (Figure 7A, Table S4). We noted that Plp1 and Mag, which were associated with central nervous system and abnormal nervous system and were also known to be located in extracellular space, interacted directly with each other. Alphafold2-Multimer^48–50^ was utilized to predict the complex structure of Mag and Plp1. It showed that the two alpha carbons of the lysine residues FSKNYQDY of Plp1 and YFNSPYPKNYPPVVF of Mag were located within 13.9 Å (Figure 7B). Since protein-protein interactions can occur if proteins exist within 20 Å of each other,^51^ it is possible that these adjacent peptides bind with each other. FSKNYQDY showed a greater decrease in accessibility in AD than in NC (Figure 7C), and YFNSPYPKNYPPVVF exhibited a similar accessibility in AD and NC at 6 months, but the lysine sites became inaccessible during AD progression, whereas no difference was observed in NC (Figure 7D).

**Figure 7.**
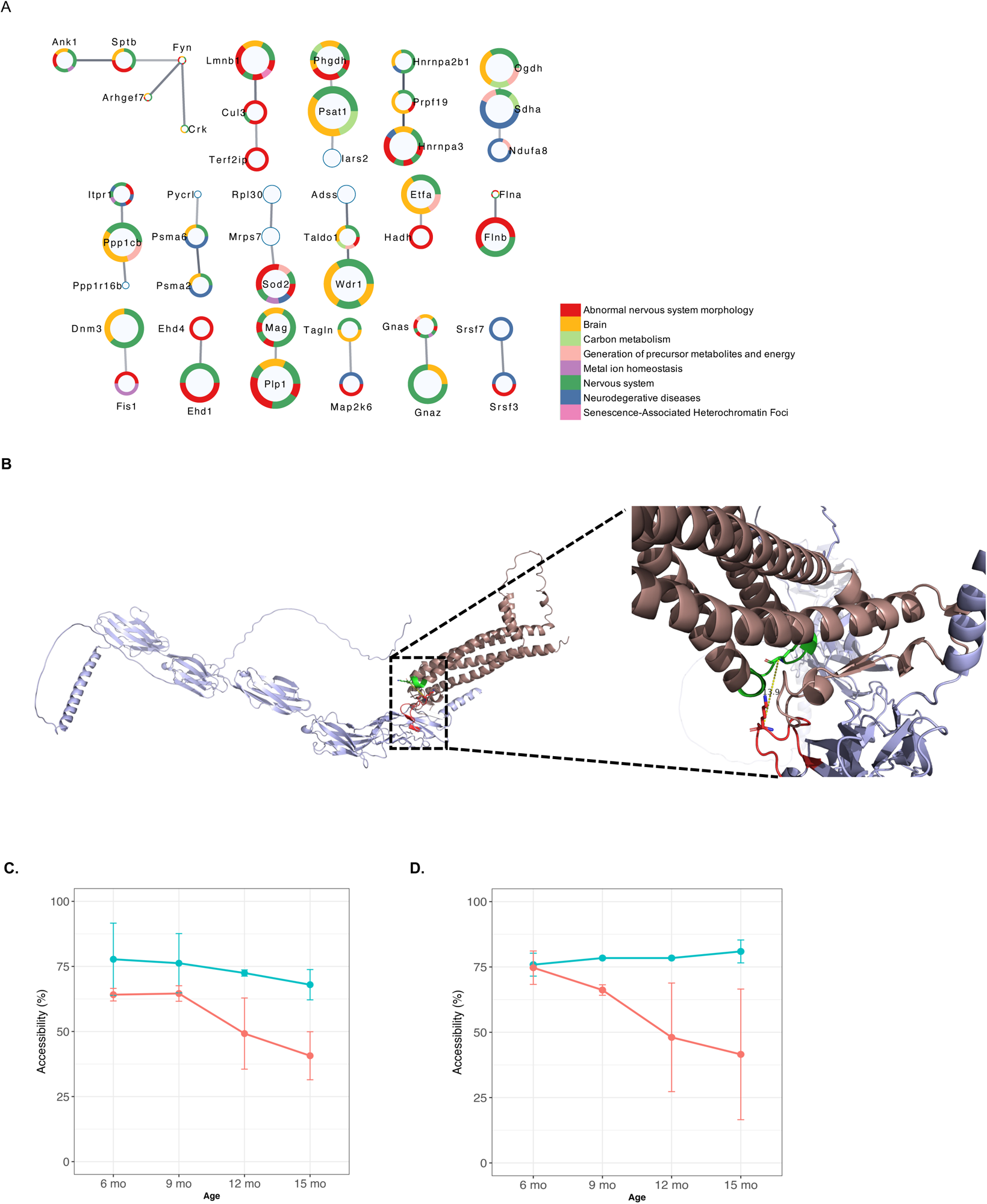
The physical interactions of the proteins in M3 of brain. **A.** Forty-five proteins in the brain dataset were physically interacted. Each node indicates a protein. The ring color of the node indicates the terms that the protein is associated with. The size of node represents the number of significantly changed lysine sites, with very small nodes indicating no significantly changed peptides, small nodes indicating one significantly changed peptide, medium nodes indicating 2-4 significantly changed peptides, and large nodes indicating more than 4 significantly changed peptides. **B.** The structure of the Plp1-Mag complex was predicted using AlphaFold-Multimer. The structure in dark pink is Plp1 and the structure in light purple is Mag. FSKNYQDY of Plp1 and YFNSPYPKNYPPVVF of Mag were presented in green and red, respectively. The right panel is an enlarged view of the complex on the left. The distance between alpha-carbons of two lysine sites was 13.9 Å. **C, D.** Structural changes in adjacent peptide regions with potential binding, with variation of the accessibility of site in AD (pink) and NC (green) for FSKNYQDY (C) and YFNSPYPKNYPPVVF (D).

In addition to the brain, it has been reported that the peripheral system plays a role in amyloid-β clearance. Approximately 40%–60% of brain-derived amyloid-β is transported across the blood-brain-barrier into the peripheral system for clearance, although the involved periphery and the mechanisms remain unclear.^52^ Spleen is composed of a variety of immune cells (with 7-8% of all cells being monocyte/macrophage) and has a role in blood filter and immunological functions. In addition, the spleen monocytes/macrophages are reported to be involved in clearing amyloid beta.^53^ Still, the physiological mechanisms underlying the association between the peripheral organs and AD remain unknown. In the ontology network of the spleen dataset, we noted “carbon metabolism” (node-147) and “neutrophil degranulation” (node-151). In these communities of proteins, the lysine sites in AD became exposed during progressing AD, while the accessibility in NC remained stable from 6 months to 15 months (Figure S11A-B). No significant change in the expression of proteins was observed for proteins of carbon metabolism (Figure S11D), but by 15 months the expression of proteins in neutrophil degranulation differed significantly from the expression at 12 months (Figure S11C). The results from the other tissues are shown in the supplementary data (Figure S12-S13 and Table S5-S8). Collectively, the results presented here provide compelling evidence of a relationship between conformational changes and protein expression, thereby highlighting the significant impact of organ-specific alterations of biological function during the progression of AD.

## Discussion

This study elucidated the AD-associated conformational changes in the proteomes of seven tissues in mice. We used the AD mouse model (APP^NL-F^), which expresses APP at wild-type levels while producing elevated pathogenic Aβ through an APP knock-in approach. This model reduces the risk of artificial phenomena that might be observed with an APP overexpressed mouse model and enhances the interpretability of the results.^21^ We found co-regulated proteins whose accessibility changed in 4 of 7 tissues, and we connected the structural protein differences in progressing AD compared to normal aging of unaffected mice to possible alterations in their biological functions. Whole animal perfusion was used to deliver reagents to comprehensively dimethyl label mouse organs with minimal intervention such as organ excision, tissue homogenization, and protein extraction that could denature or alter proteins. Our quantitative method for accessibility measurement determines the relative fraction of inaccessible over accessible for each lysine site. Changes in accessibility can be interpreted as a change in protein folding or a change in interaction with another molecule and thus can be a surrogate for protein conformation changes between different conditions.

In this study, we focused on changes in proteins of the brain. Amyloid beta is a well-known protein that accumulates as plaques in the brain and serves as a marker of AD, but we were not able to detect structural changes of amyloid beta. We quantified the accessibility of three peptides from APP in only one 15-month mouse. It has been known that the amyloid beta plaque accumulates primarily in the cortex, followed by the hippocampus, basal ganglia, thalamus, and basal forebrain.^54^ However, because the purpose of this study was to globally investigate the change of the proteome, we used whole brain tissue sample in this study, not just the amyloid-beta plaque containing cortex region of the brain.

It has been suggested that kidney function is linked to brain activity,^55^ and changes in kidney function may play a role in the development and progression of AD. Studies have shown that the MRI images of the brains of AD patients were similar to that of kidney patients^56^ and a systematic meta-analysis demonstrated that cognitive impairment is significantly related to malfunction of kidney.^57^ Despite persuasive evidence of the link between the kidney and AD, the exact physiological mechanisms underlying this relationship are not fully understood. From our WGCNA analysis, we found two modules (M11 and M14) to be significantly co-expressed in kidney and investigated the enriched functions of the structurally altered proteins using GO enrichment analysis. In both modules, purine-related functions were most significantly enriched. Our findings from kidney are supported by results of metabolomic studies showing that guanosine monophosphate (which is derived from purine guanine and associated with purine metabolism) was dysregulated in the brain of an AD mouse model based on APOE4 allele mutant mice.^58^ Therefore, it can be inferred that purine metabolism may play a role in the link between kidney function and AD.

We introduced a feature to the CPP pipeline that can be employed under various experimental conditions. A multidimensional protein identification technology (MudPIT) can be used to generate high sequence coverage of the proteome.^59, 60^ MudPIT separates peptides first by charge and then by hydrophobicity to create a two-dimensional separation. In MudPIT, peptides released from a C18 trap column are loaded onto a strong cation exchange column and then released with a buffer that increases in ionic strength through sequential elution steps. Released peptides are bound to the C18 analytical column and are then eluted sequentially to the mass spectrometer based on hydrophobicity. It is not recommended that hydrophobicity be used for the first dimension of the 2D-LC-MS because the hydrophobicity of deuterium is slightly lower than hydrogen, so a partial separation of differentially labeled peptides might occur.^61^

Limitations to this study include the lack of a standard to monitor the distribution of reagent solution to organs in the body during the first labeling step. Although some signs such as body twitching, tail flicking, and head moving in the anesthetized animals was observed, a reliable quantitative standard to assess the extent of labeling in each organ would be useful. We defined labeling efficiency as the ratio of initially labeled peptides over identified lysine-containing peptides per tissue. It is an inevitable limitation of CPP that dimethylation cannot occur on a lysine site that is already modified (i.e., acetylation and ubiquitination); thus, the accessibility of innate partially labeled lysine sites will not be accurate. An additional limitation is the method uses lysine as the conformation reporter which occurs at a frequency of roughly 5-7% in proteins and thus the quality of the analysis scales with sequence coverage of individual proteins. High sequence coverage will generate more MS/MS of lysine containing peptides and increase the completeness of the analysis.

It is likely that changes in protein structure are a result of failing proteostasis and expression changes are a function of alterations in protein synthesis and/or degradation. Mutations of DNA in somatic cells accumulate as we age ^62–65^ and can result in changes to protein sequences, necessitating more effort to keep proteins properly folded. Our new method provides a means to measure protein surface accessibility as a surrogate for protein conformation *in vivo* and in animal models to study of the role of protein folding in aging and AD. Our proteomic analysis showed changes in protein structures in multiple tissues during the progression of AD. Even though the patterns of change in non-brain tissue did not correspond exactly with those in the brain tissue (and it is not clear they should), our analysis showed changes in protein structure and expression in other tissues in the AD mouse model. In conclusion, this new method to measure *in vivo* alterations to protein surface accessibility in animal models of disease provides a means to measure a previously unexplored characteristic of proteins to provide insights into how physiological systems are perturbed.

### Data availability

All mass spectrometry raw data in this study have been deposited to MassIVE repository with identifier MSV000091970.

## Acknowledgements

We thank Dr. Claire Delahunty for critical reading the manuscript. This work was supported by NIH/NIA grant RF1AG061846-01 and 5R01AG075862.

## Author contributions

A.S., C.B. and J.R.Y. conceived the project. A.S. and H.K. performed the experiments. J.D. measured the samples on the mass spectrometer. A.S and H.K. analyzed the data and the results. C.B. and D.M. provided the critical feedback. J.R.Y. supervised the project. A.S. wrote the manuscript and prepared the figures with help from all authors.

## Declaration of interests

The authors declare no competing interests.

---

**Figure S1.**
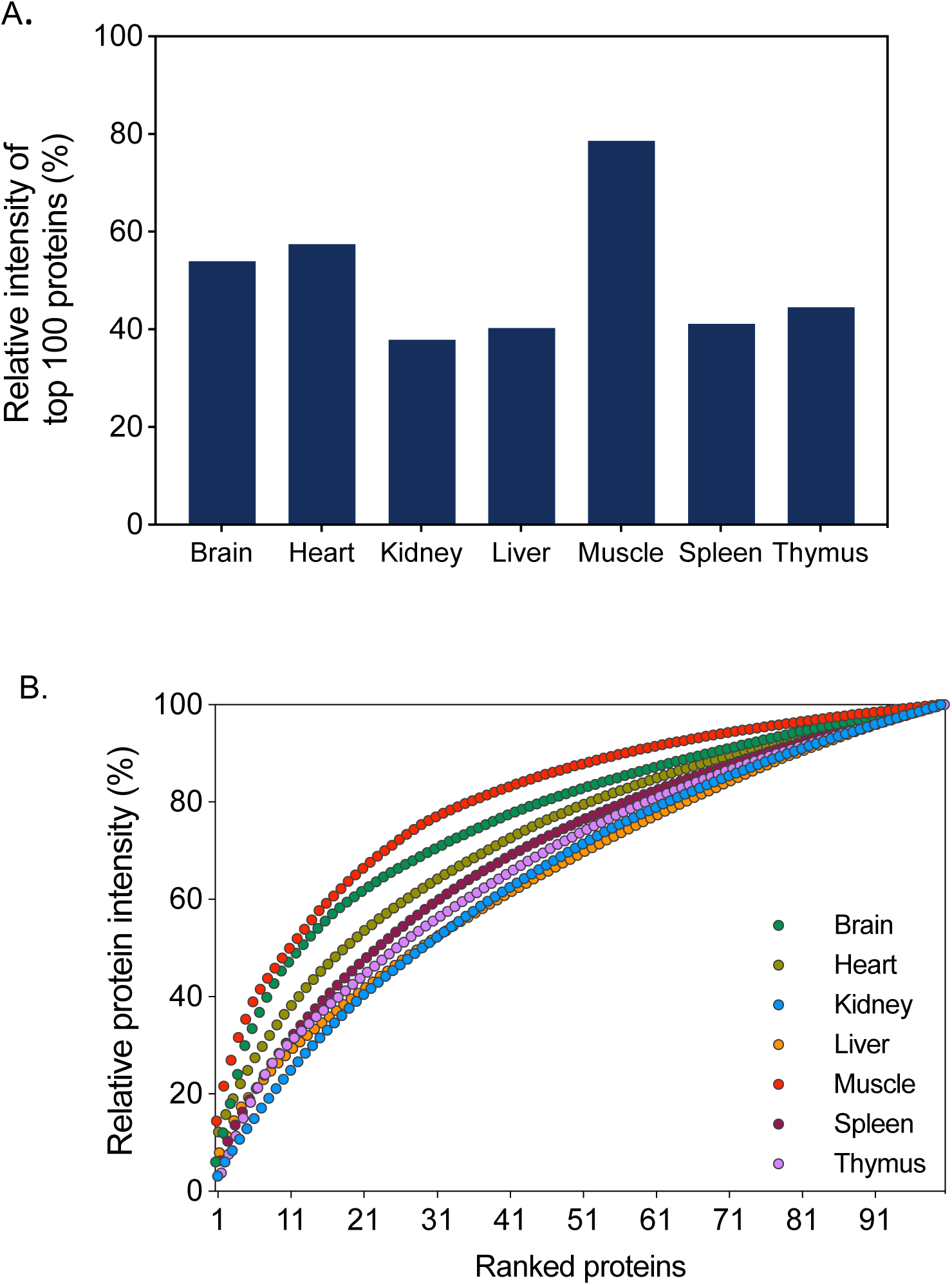
Abundance of the 100 most abundant proteins in each tissue. (A) Relative abundance of the 100 most abundant proteins in each tissue. Abundance calculation was based on protein intensity relative to the overall intensity. The sum of intensity of the top100 proteins contributed over 78.6% to the total intensity in muscle tissue. (B) Accumulation of top 100-protein abundance in the mouse tissues. Accumulation of protein mass are plotted for 7 tissues. Proteins are ranked by their intensity.

**Figure S2.**
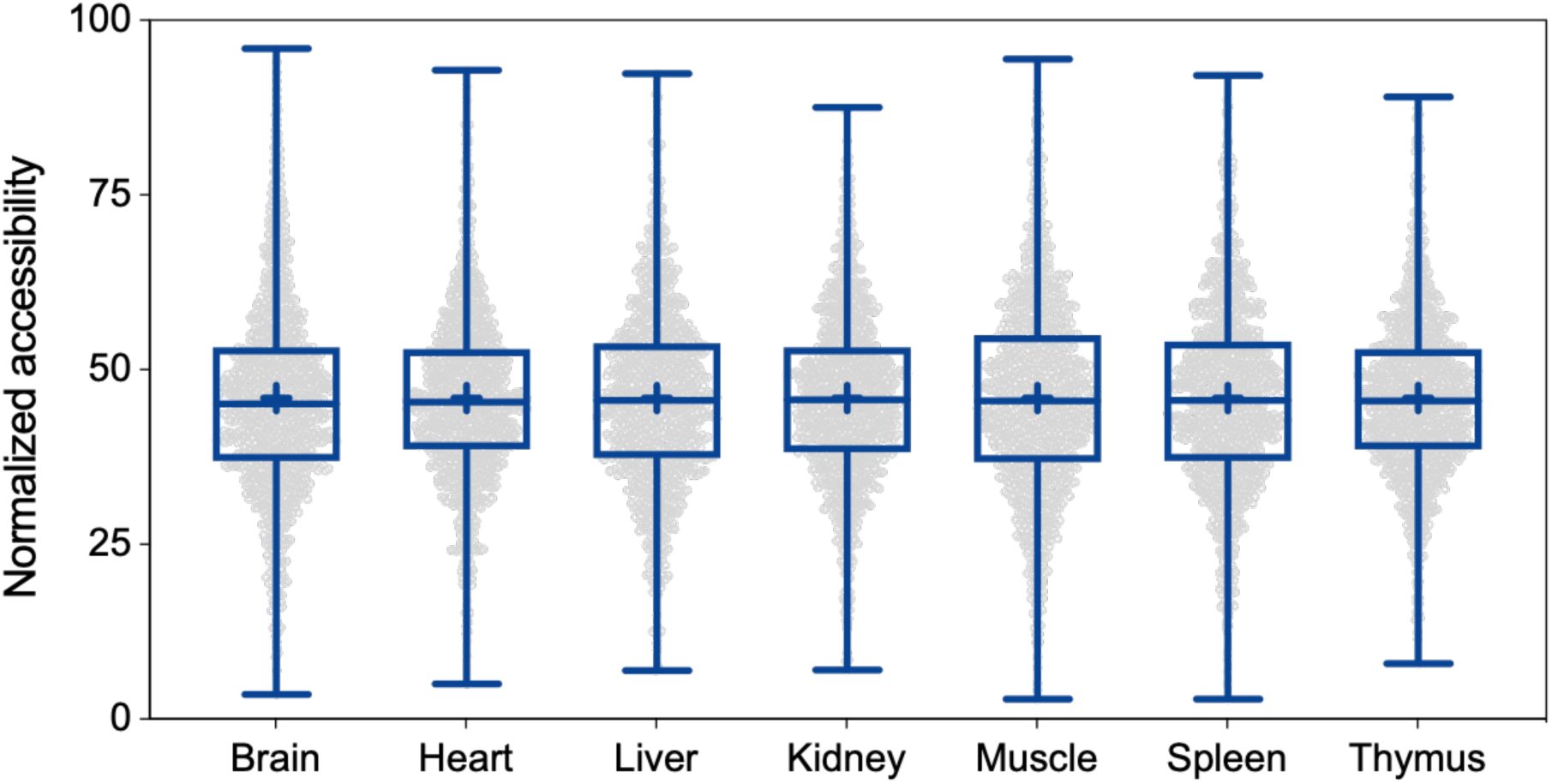
Normalized accessibility. The accessibility values of the peptides that were detected from all 7 tissues were quantile-normalized to identify the distribution of the values across 7 tissues.

**Figure S3.**
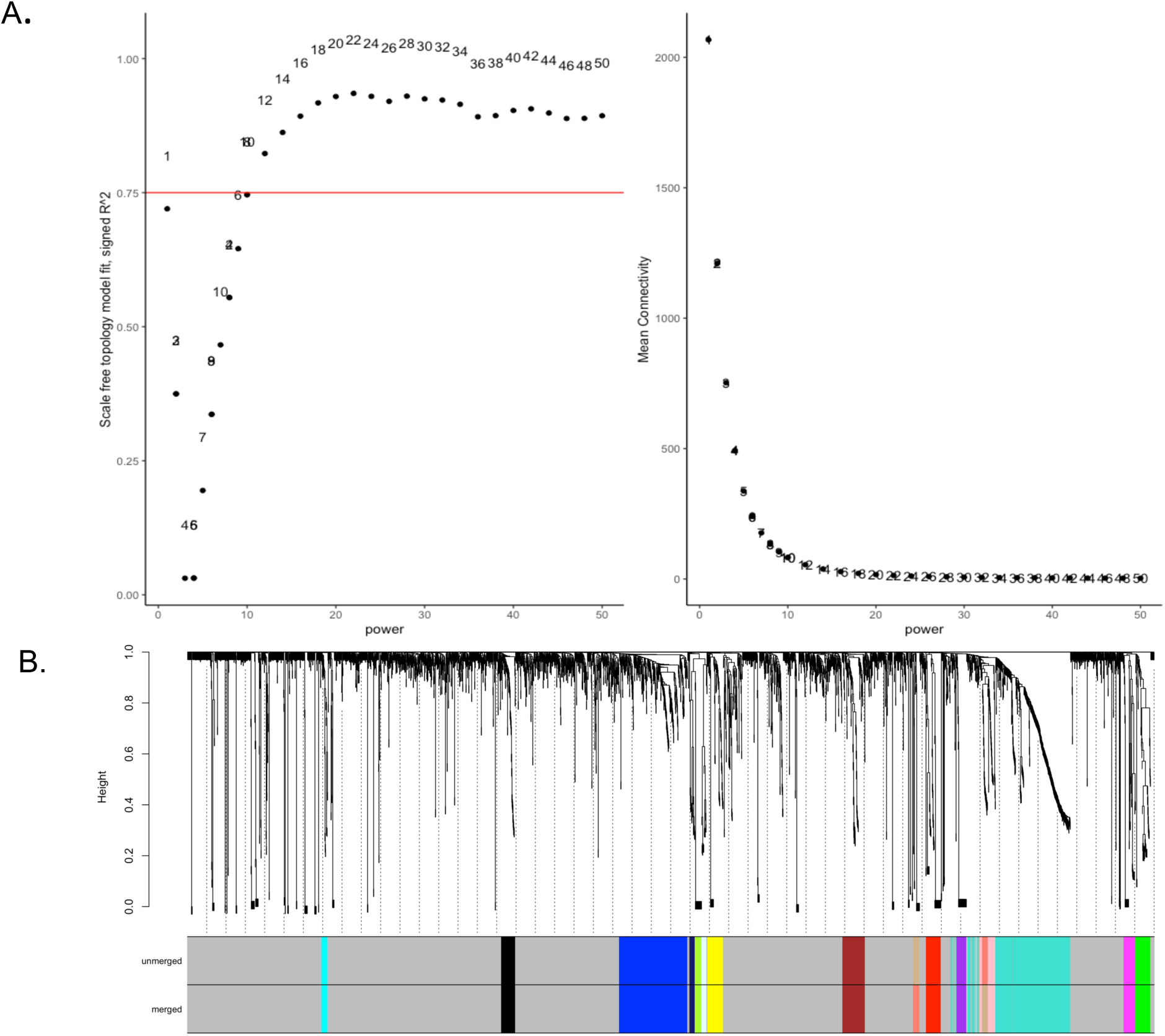
Modularization of protein co-expression. A brain dataset shown representatively. Soft threshold power was determined using the pickSoftThreshold() function in the WGCNA package. The threshold value was selected when R^2^ reaches a plateau over 0.75, indicating that the network has achieved a scale-free topology (A). The cluster dendrogram (upper) and the co-expression modules (lower) were generated by hierarchical clustering (B). The branches of the dendrogram represent individual proteins. The height indicates the Euclidean distance. Each module that contains weighted co-expressed proteins is displayed with a distinct color.

**Figure S4.**
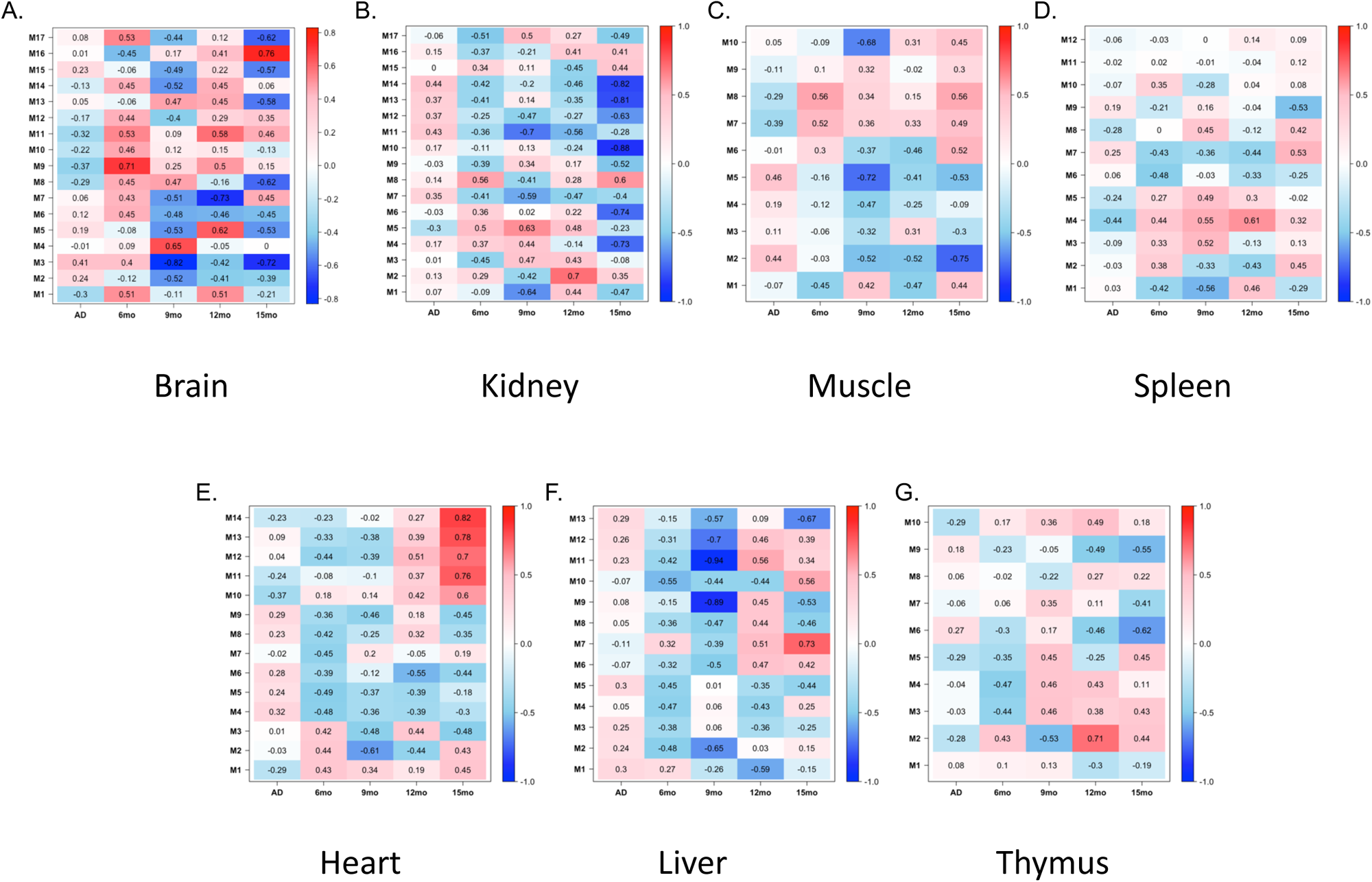
Relationship between module and traits in brain (A), kidney (B), muscle (C), spleen (D), heart (E), liver (F), and thymus (G). In the first column of each heatmap, the correlation was calculated with all samples, and in the next four columns, the correlations were calculated with samples sub-divided by age.

**Figure S5.**
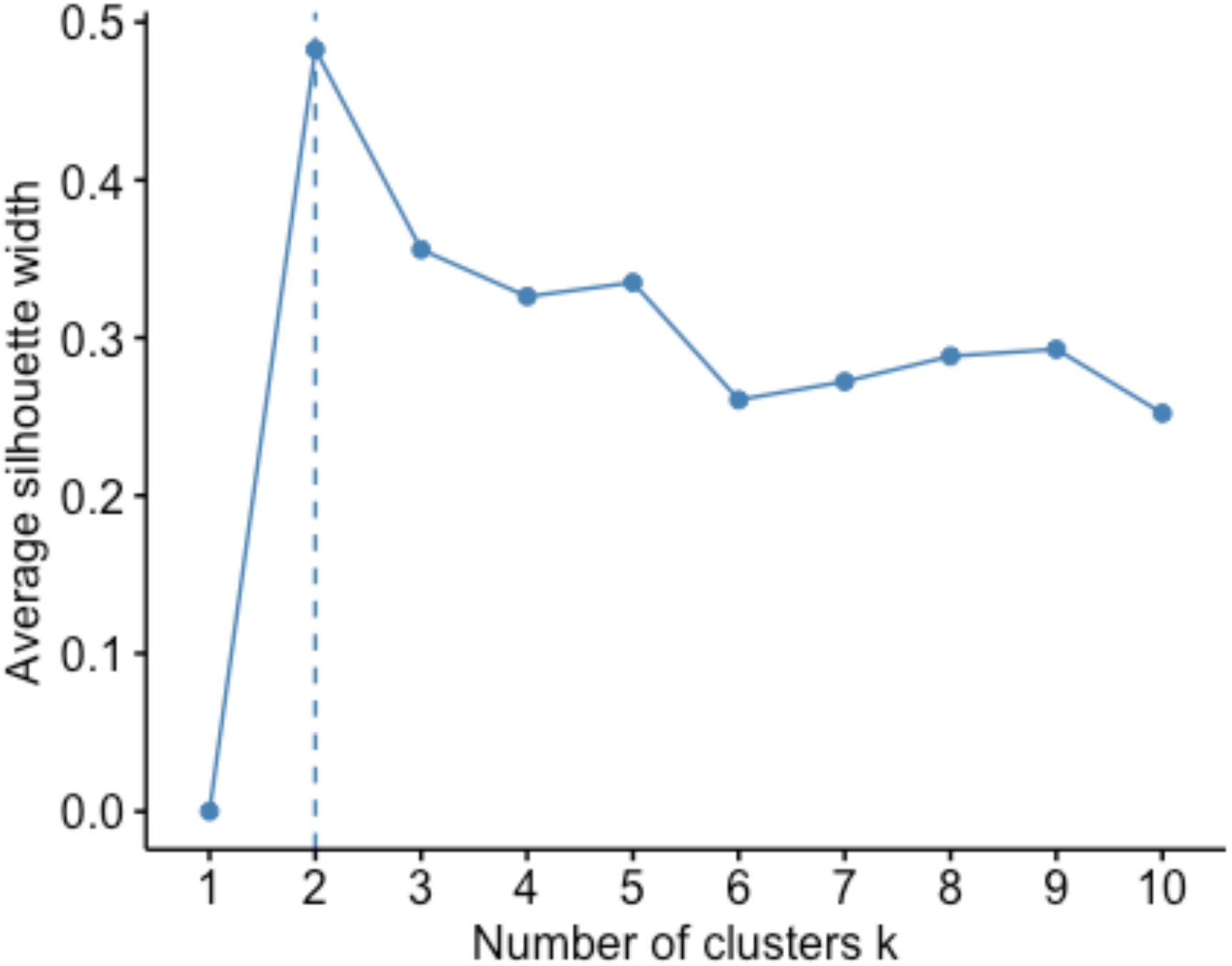
Optimization of the number of clusters in unsupervised machine learning algorithms for the brain dataset. The number of clusters was determined in K-means clustering by evaluating the quality of the clustering results for different values of k (the number of clusters). The silhouette score (silhouette width) measures how similar the data is to its own cluster compared to other clusters. A score close to 1 indicates that the data is well-matched to its own cluster. The k value that showed the highest score was selected per each tissue dataset and was determined to be 2 in the brain dataset.

**Figure S6.**
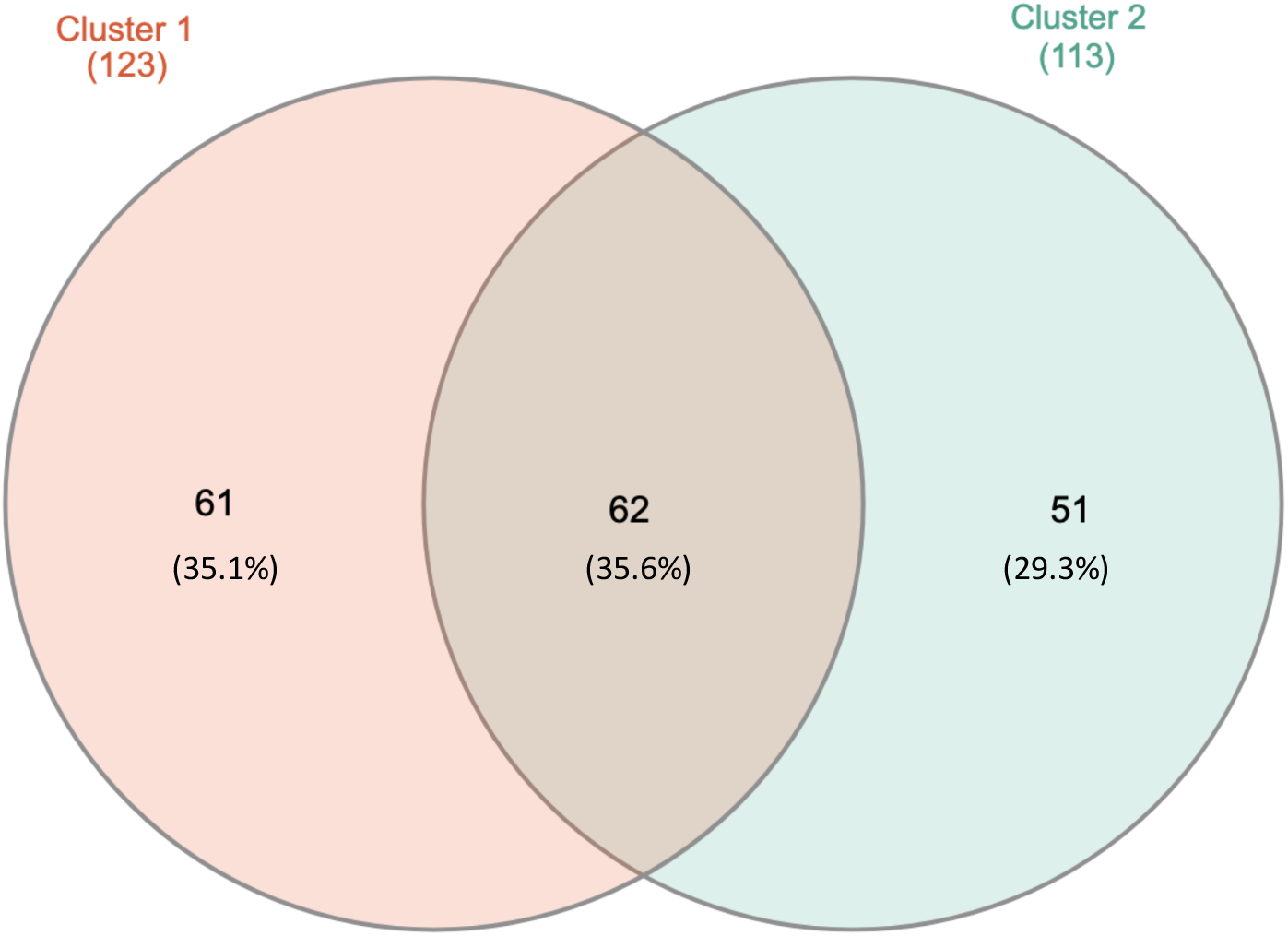
Overlapped proteins between two clusters in module 3 (M3) of brain. One hundred seventy-four of the 289 proteins of M3 were labeled with dimethylation, and the labeled peptides were clustered into two clusters. All the labeled peptides of 61 proteins were clustered into cluster 1; all the labeled peptides of 51 proteins were clustered into cluster 2.

**Figure S7.**
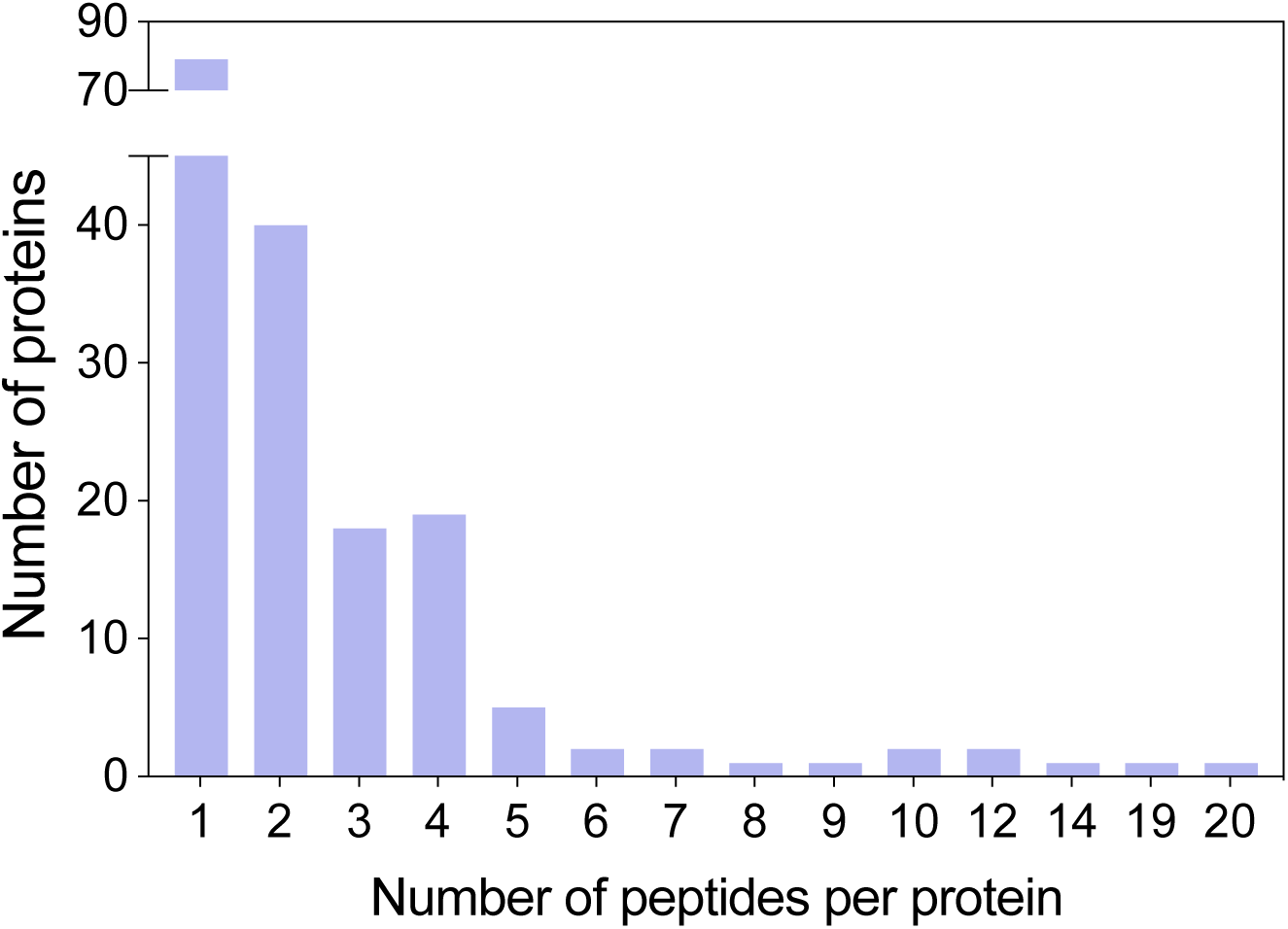
Frequency of labeled peptides per proteins in M3. Map1a showed the highest number of the labeled peptides (n = 20) per protein. Pkg1 and Mdh1 each contained 19 labeled peptides. 79 proteins included a single labeled peptide.

**Figure S8.**
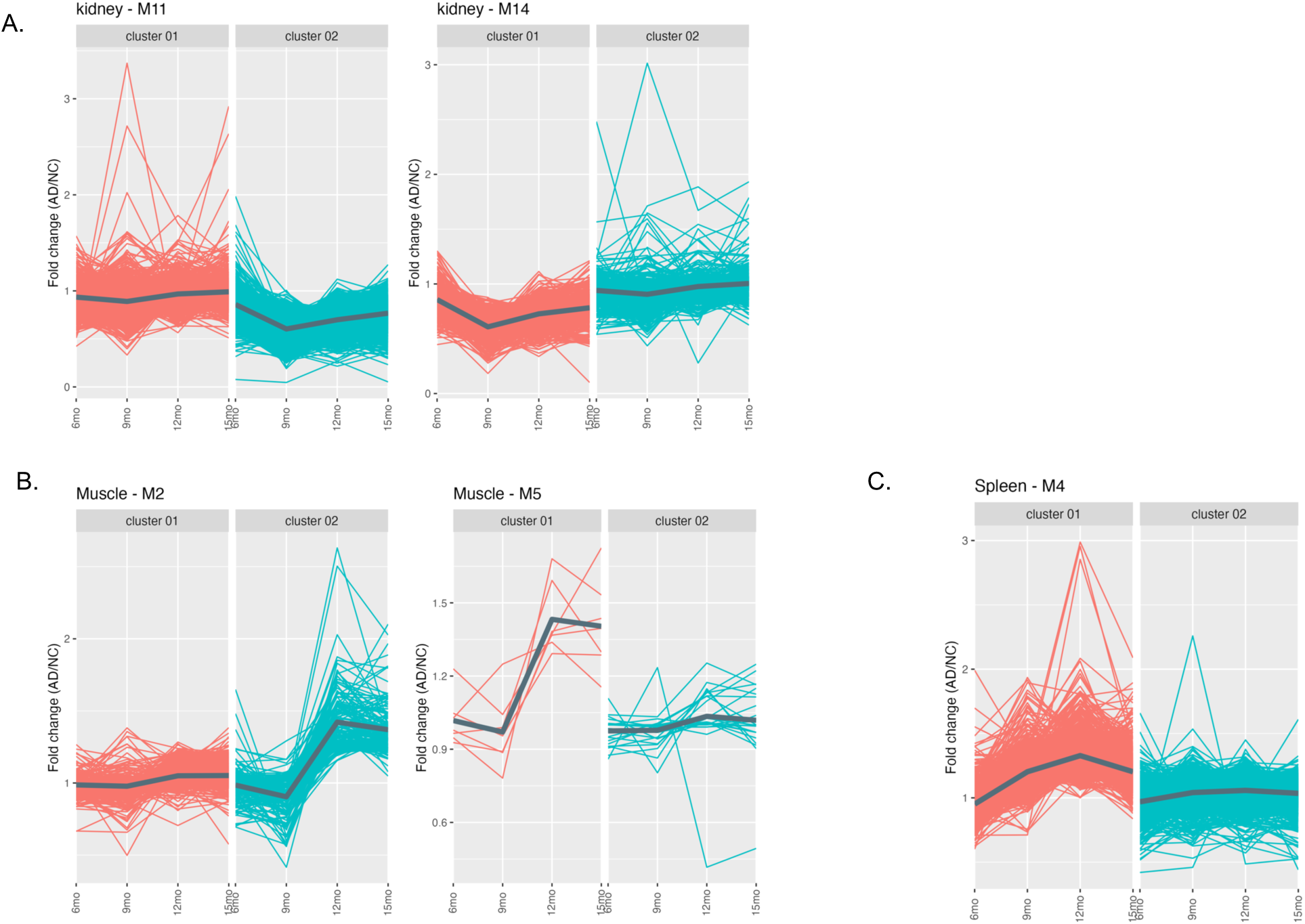
Clusters of the labeled peptides based on the fold-change of the accessibility for Kidney (A), Muscle (B), and Spleen (C). In the results of WGCNA analysis, two modules of kidney (M11 and M14), muscle (M2 and M5) and one module (M4) of spleen showed significant correlations with AD trait. Following that, the labeled peptides mapped to proteins in those modules were clustered by K-means clustering. For muscle, only 32 labeled peptides that mapped to 17 proteins were included in module 5 (M5), 7 peptides showed alterations in the fold-change of accessibility. GO enrichment analysis was not conducted with proteins of M5 due to the small number of peptides to be analyzed.

**Figure S9.**
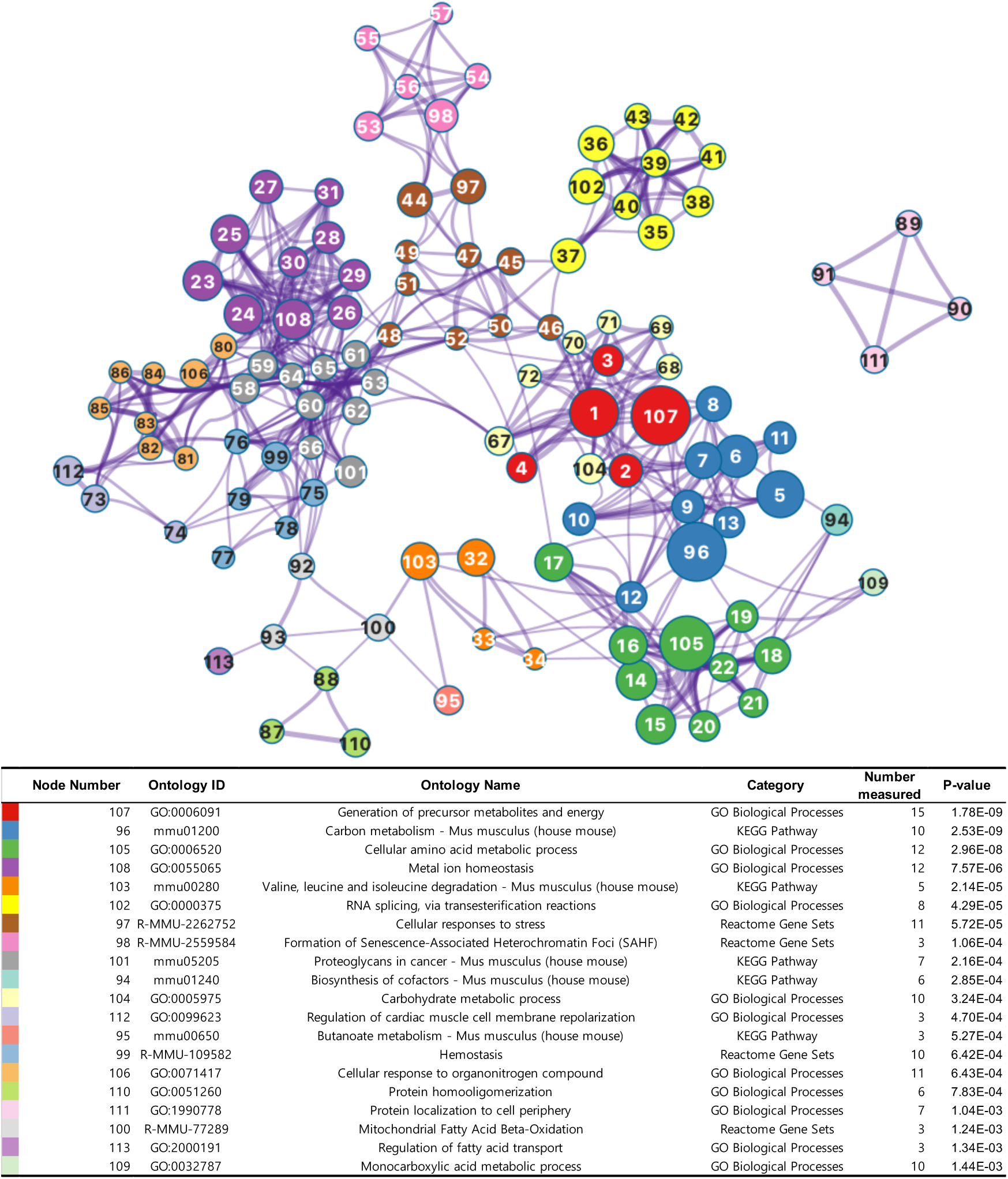
Ontology network of brain constructed by structurally changing proteins, and representative terms. Ontology networks were constructed using the proteins that harbored changed peptides in accessibility (cluster 1 of K-means clustering). Each node indicates the individual terms, and the node size represents the significance. All terms with the *P*-value less than 0.5 were enriched. The color reflects the membership of the clusters. The lower table presents the details of the representative nodes for each cluster.

**Figure S10.**
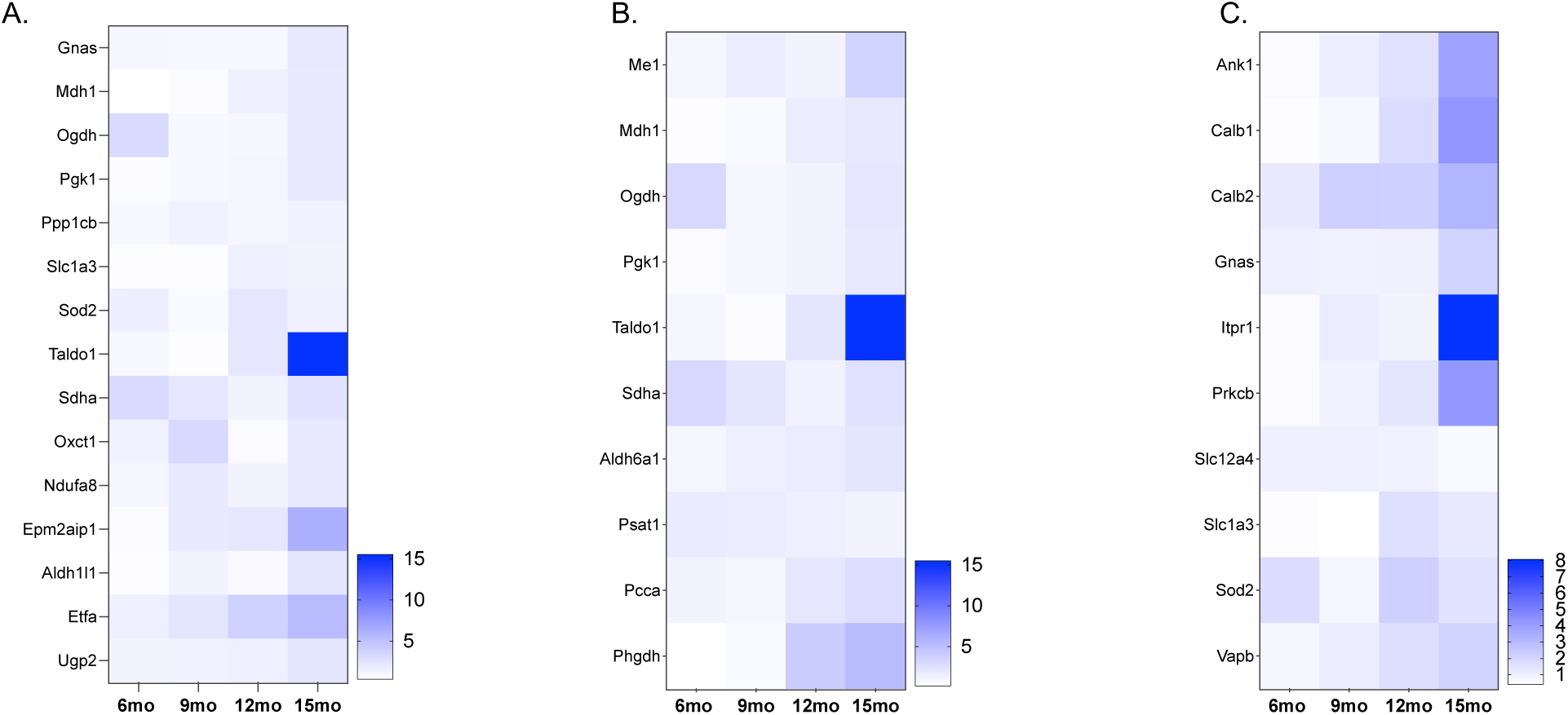
Altered abundance of proteins. Heatmap were plotted with the fold-change of the abundance, which is the value of protein abundance in AD divided by that in NC to correct for an age effect; node 107 (A), node 96 (B), and node 108 (C).

**Figure S11.**
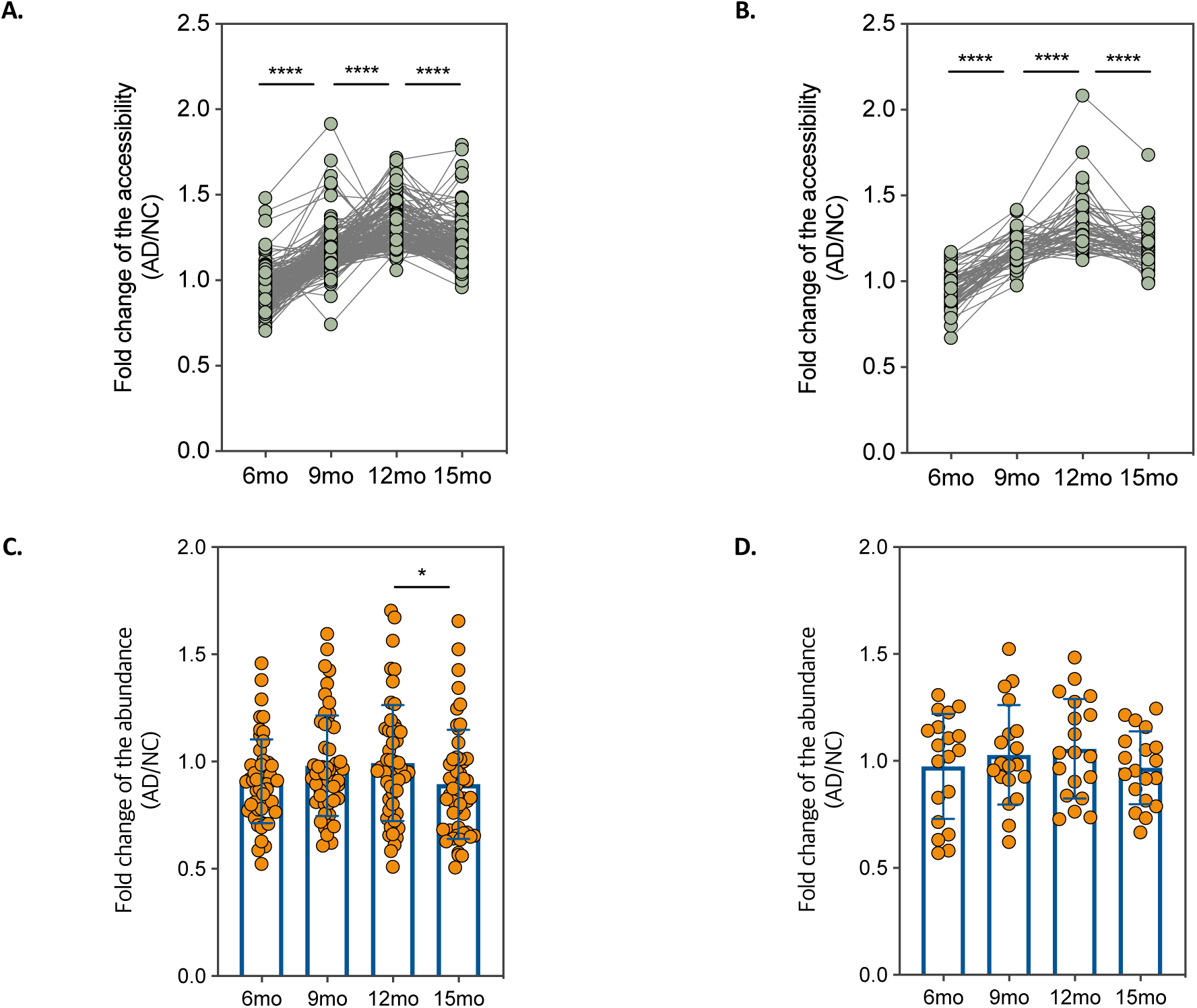
Structural changes that precede expression changes in spleen. Fold changes of the accessibility (A, B) and fold change of the protein abundance (C, D) for neutrophil degranulation (A, C) and carbon metabolism (B, D). *p < 0.05, ****p < 0.0001

**Figure S12.**
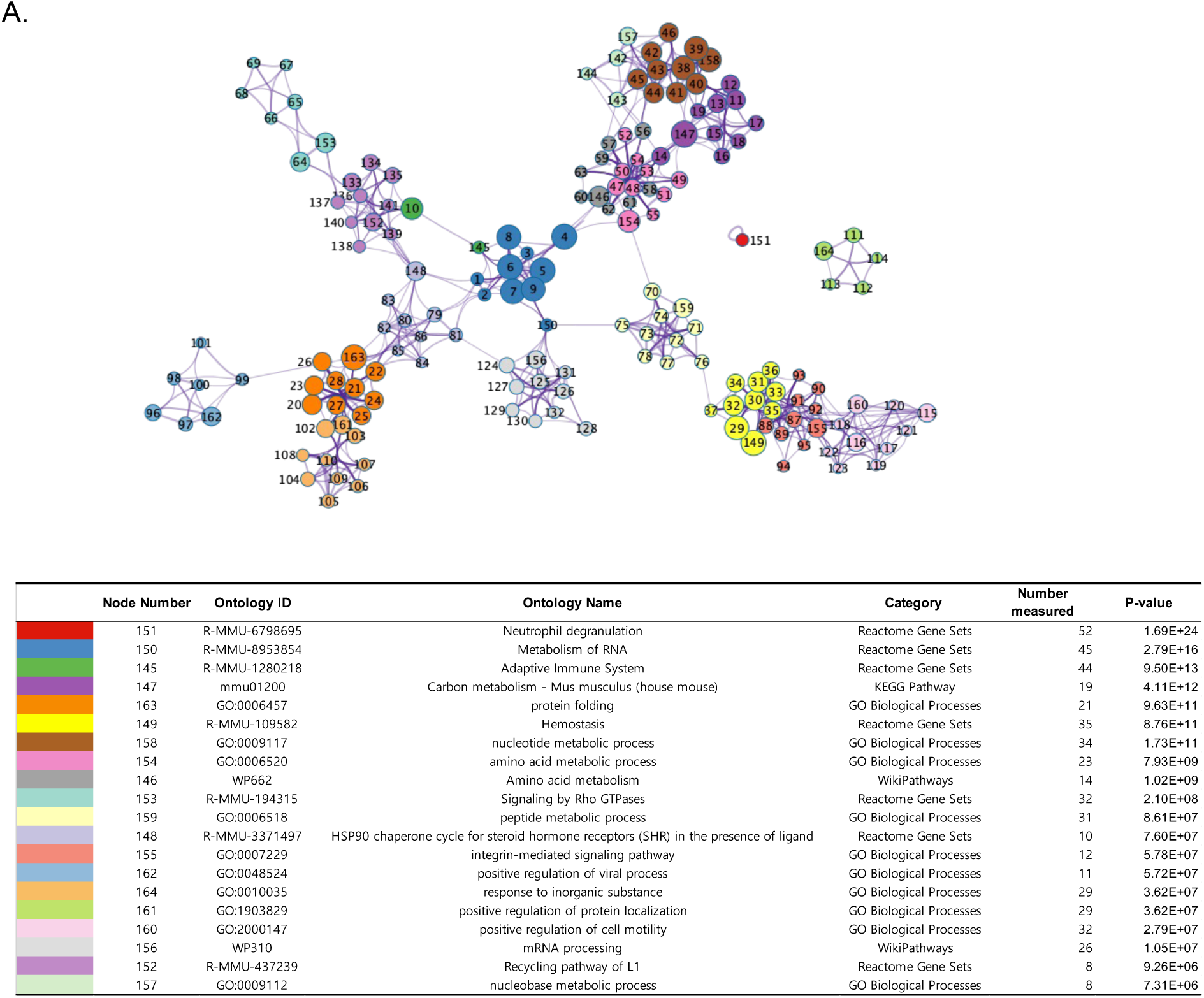
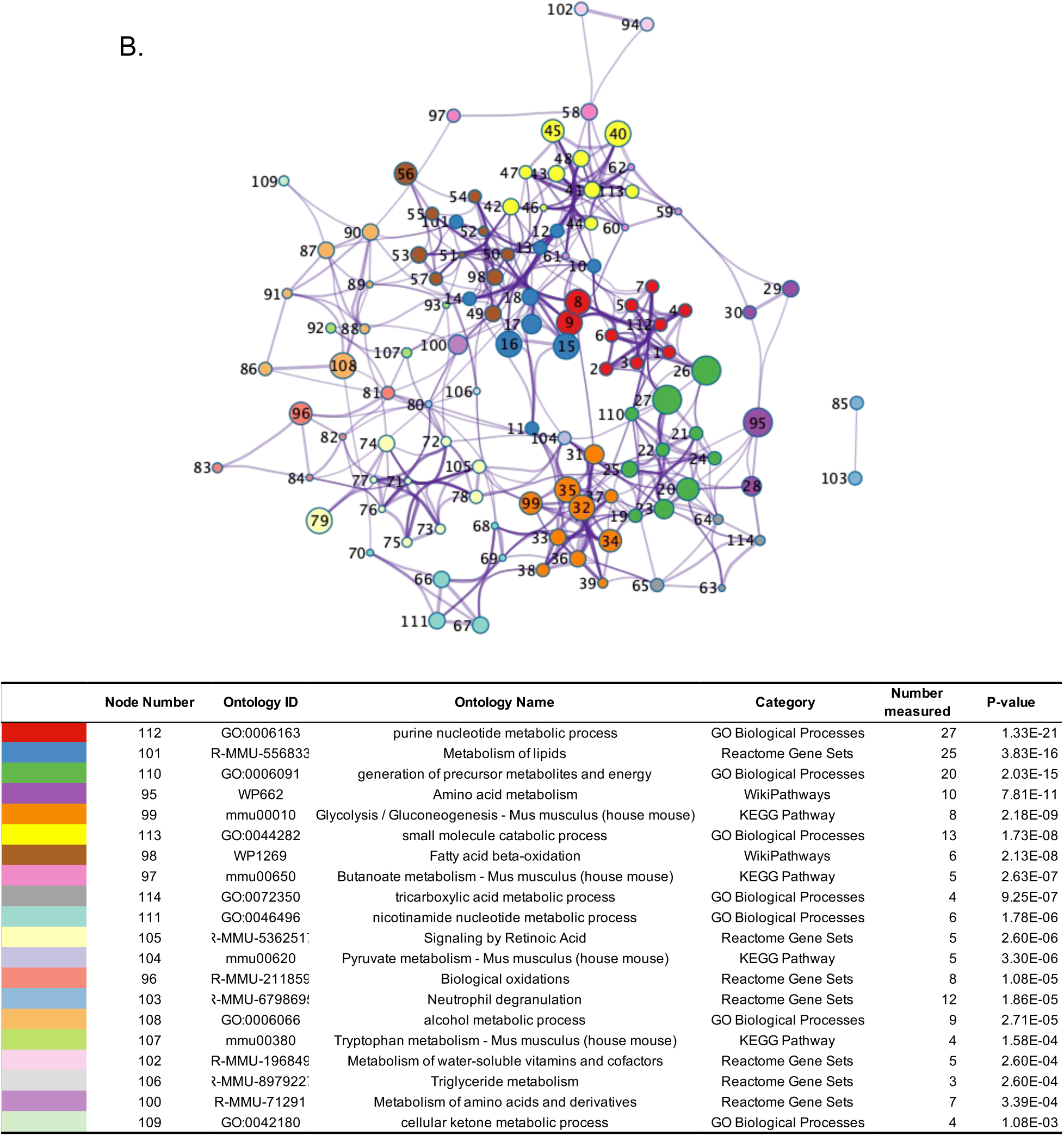
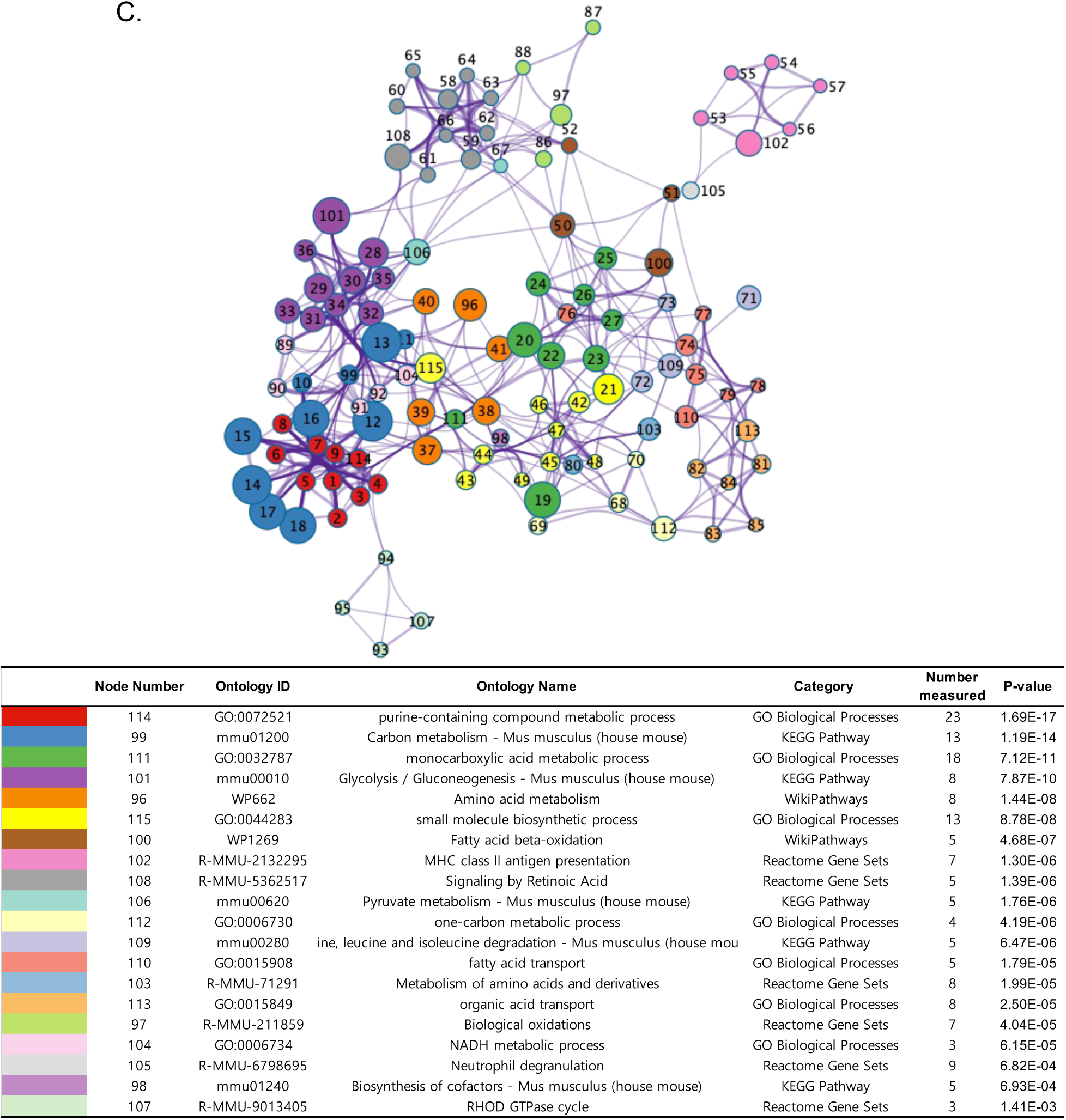
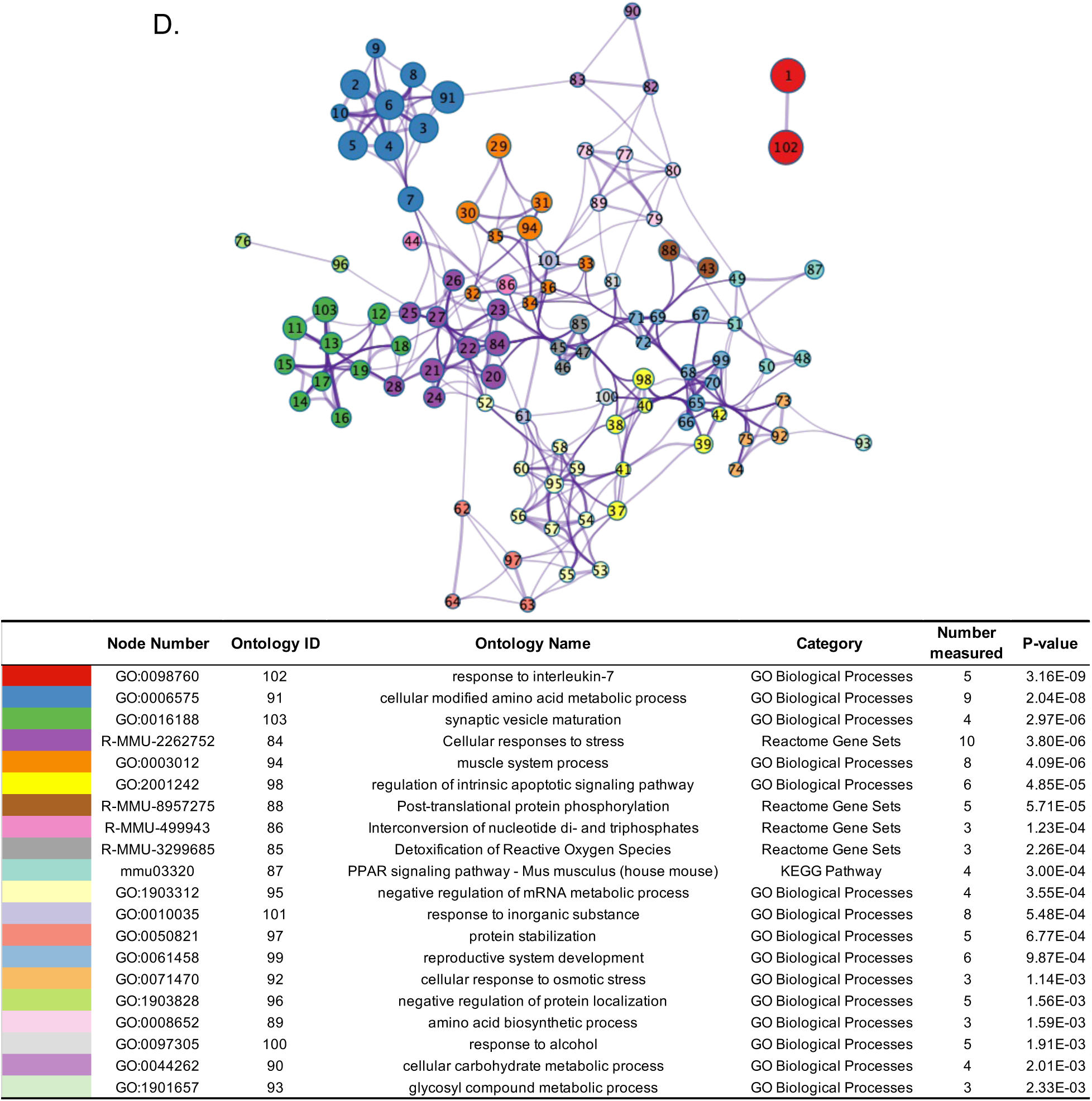
Ontology network of structural changes proteins per each dataset for spleen (A), M11 of kidney (B), M13 of kidney (C), and muscle (D).

**Figure S13.**
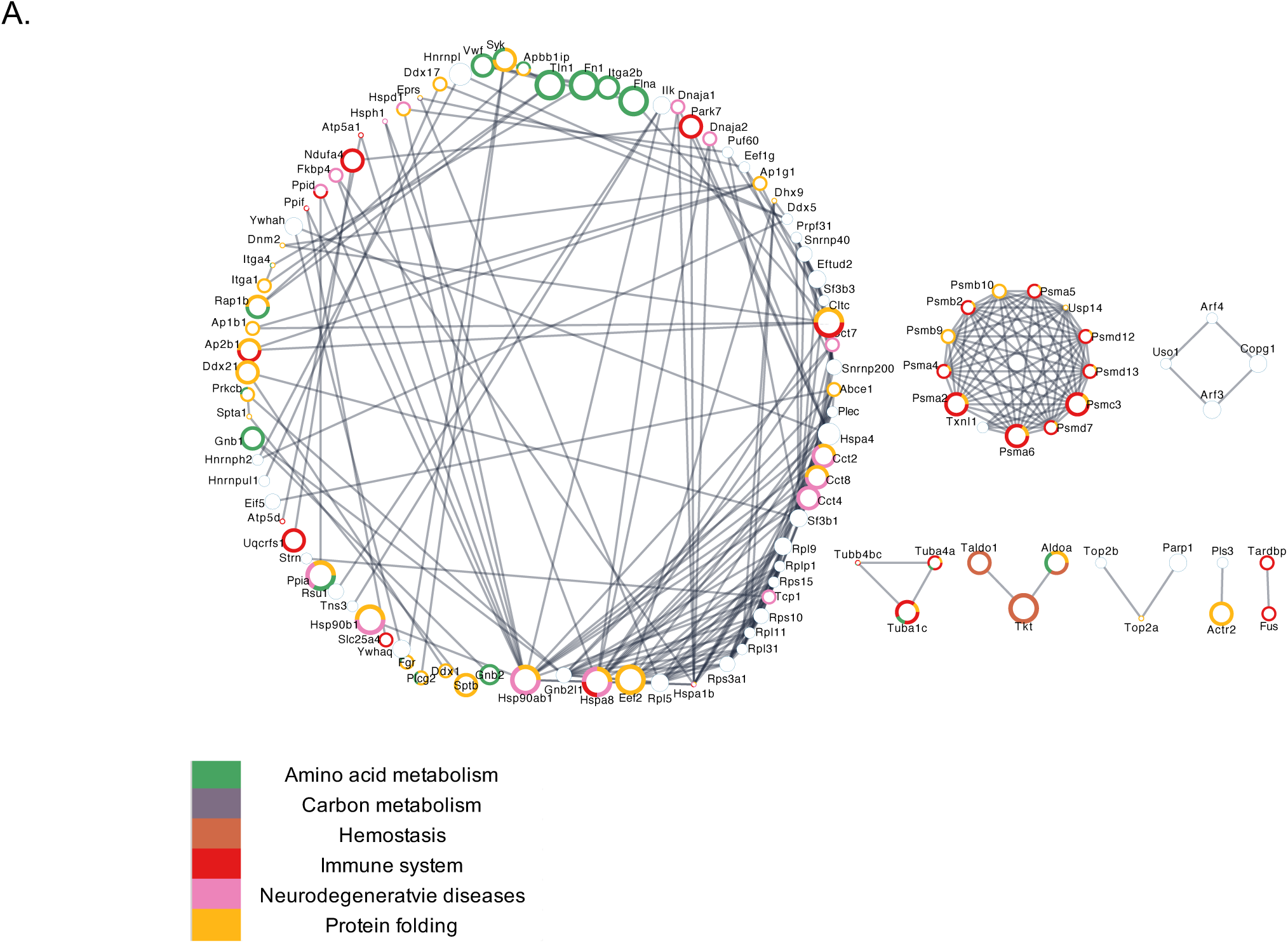
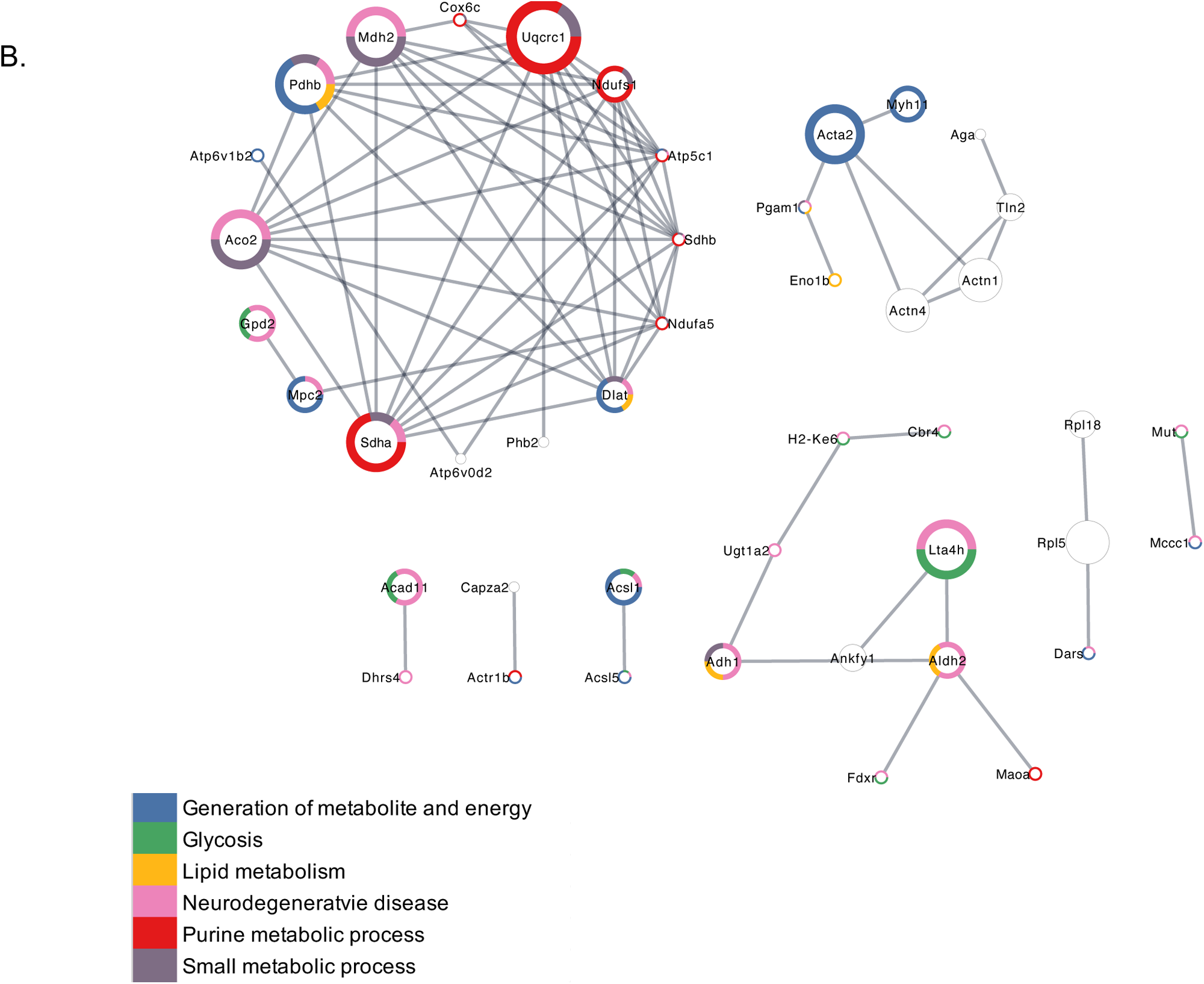
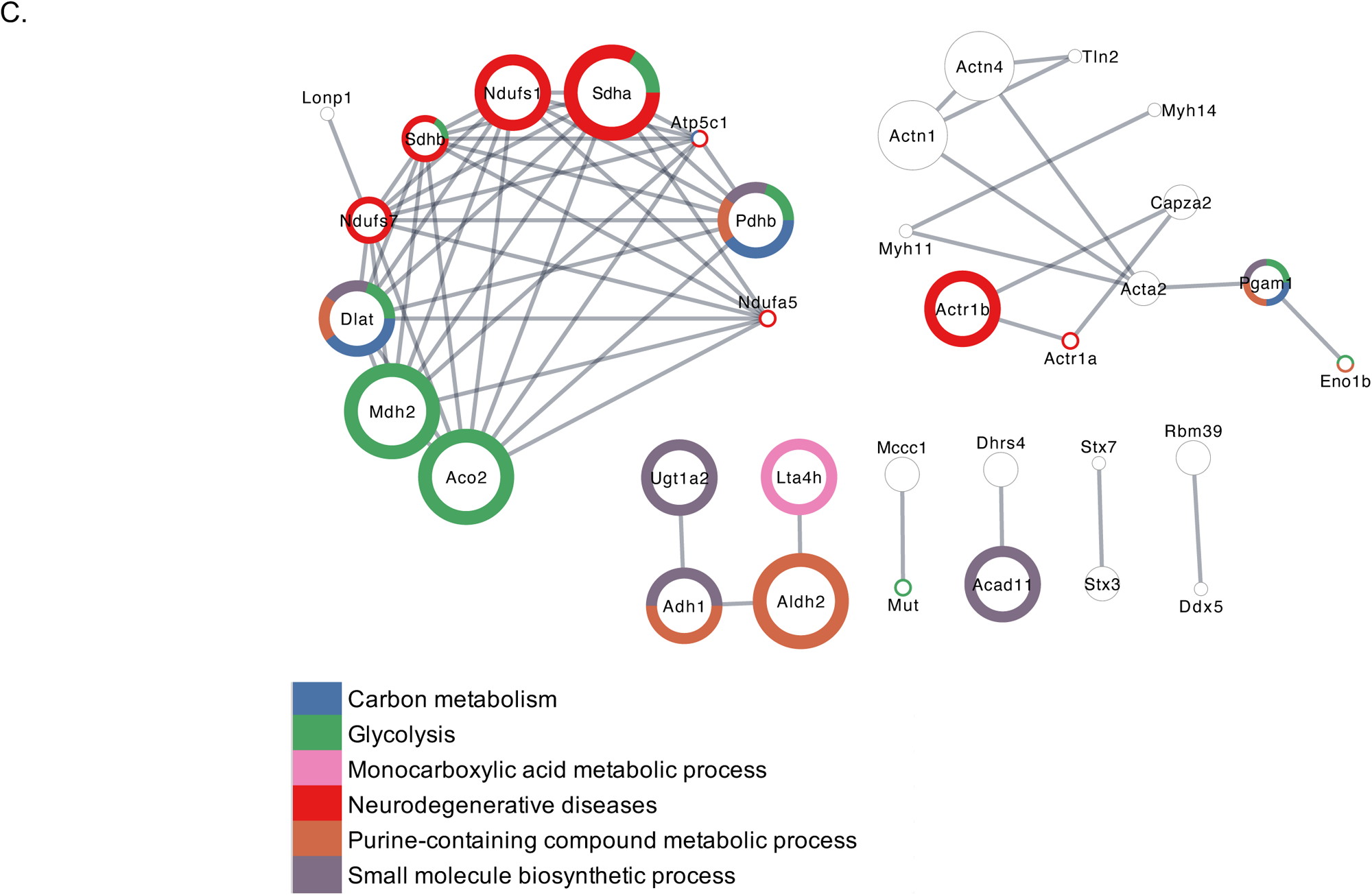
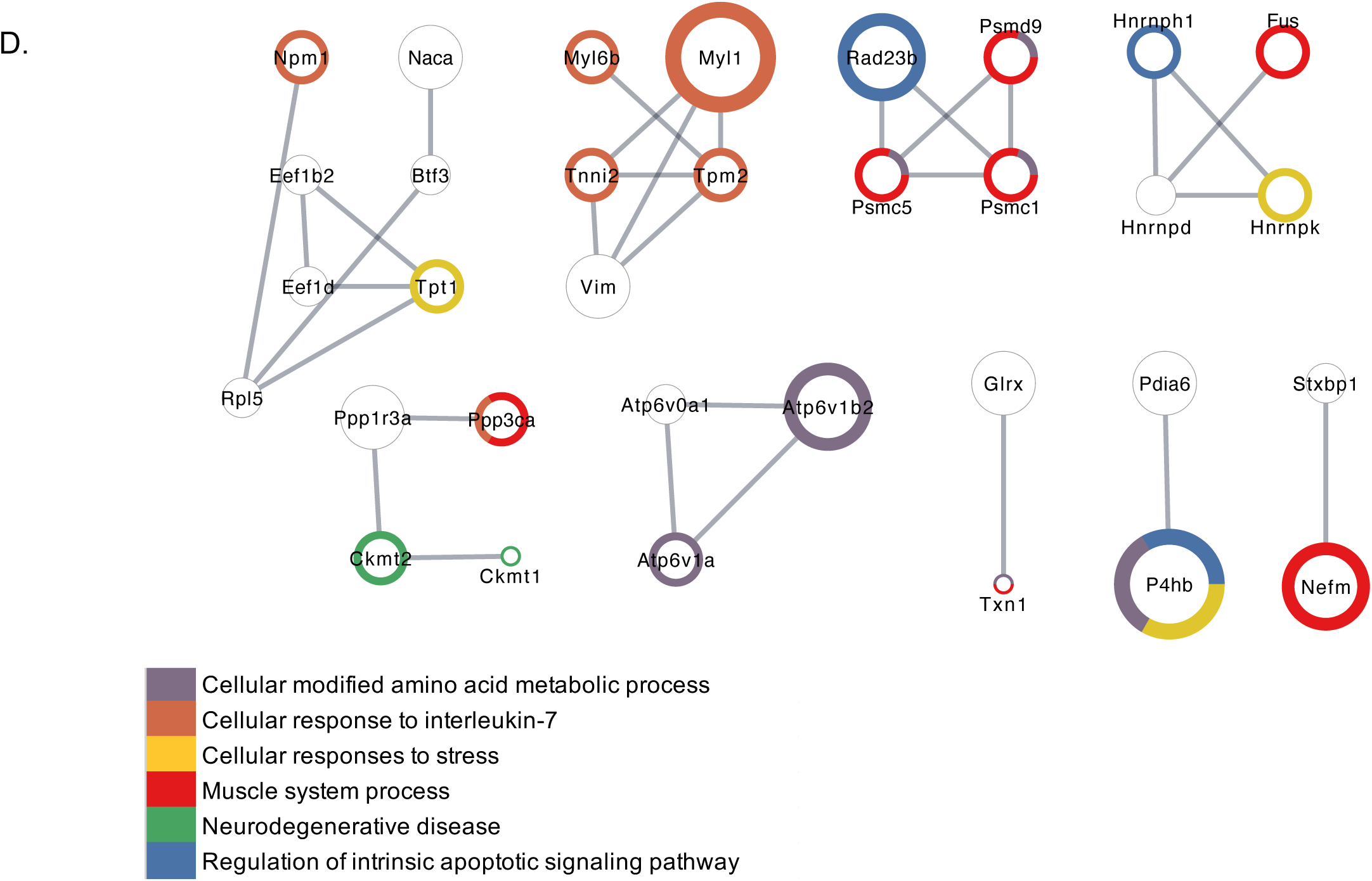
Physical interactions of proteins and their associated functions for spleen (A), M11 of kidney (B), M13 of kidney (C), and muscle (D). The ring color of the node indicates the terms that the protein is associated with. The size of node represents the number of significantly changed lysine sites.

## Notes

### Competing Interest Statement

The authors have declared no competing interest.

